# Joint Motion Estimation and Source Identification using Convective Regularisation with an Application to the Analysis of Laser Nanoablations

**DOI:** 10.1101/686261

**Authors:** Lukas F. Lang, Nilankur Dutta, Elena Scarpa, Bénédicte Sanson, Carola-Bibiane Schönlieb, Jocelyn Étienne

## Abstract

We propose a variational method for joint motion estimation and source identification in one-dimensional image sequences. The problem is motivated by fluorescence microscopy data of laser nanoablations of cell membranes in live Drosophila embryos, which can be conveniently—and without loss of significant information—represented in space-time plots, so called kymographs. Based on mechanical models of tissue formation, we propose a variational formulation that is based on the nonhomogenous continuity equation and investigate the solution of this ill-posed inverse problem using convective regularisation. We show existence of a minimiser of the minimisation problem, derive the associated Euler–Lagrange equations, and numerically solve them using a finite element discretisation together with Newton’s method. Based on synthetic data, we demonstrate that source estimation can be crucial whenever signal variations can not be explained by advection alone. Furthermore, we perform an extensive evaluation and comparison of various models, including standard optical flow, based on manually annotated kymographs that measure velocities of visible features. Finally, we present results for data generated by a mechanical model of tissue formation and demonstrate that our approach reliably estimates both a velocity and a source.

## 1 Introduction

### 1.1 Motivation

Motion estimation is a ubiquitous and fundamental problem in image analysis, see e.g. [5]. It is concerned with the efficient and accurate estimation of displacement fields in spatio-temporal data and has a wide range of applications, not necessarily limited to natural images. Optical flow [30] is one popular example of motion estimation, which designates the apparent motion of brightness patterns in a sequence of images and is based on the assumption of constant brightness. Recently, optical flow methods have been used for the quantitative analysis of biological image sequences on cellular and subcellular level. See, for instance, [2, 9, 10, 21, 31, 34, 39, 44, 50]. While the concept is in practice well-suited for natural scenes, the use of the less restrictive continuity equation, which arises from mass conservation, can be more favourable in certain scenarios. For instance, in [15, 16] it is used for fluid flow estimation.

In developmental biology, the study of the morphogenesis of model organisms is specifically calling for image analysis methods that are able to extract time-dependent deformations and flow velocities from microscopy image sequences. Morphogenesis is the process that leads to an organism developing its shape as a result of the implementation of a genetic programme [28] and includes, among other mechanisms, tissue deformations. These deformations can be observed through video microscopy by fluorescently labelling molecules that are associated with compounds of mechanical relevance within an embryo [32].

In many cases, these molecules are organised spatially in discrete structures, such as cell membranes. To compute deformations on the level of these molecular structures, segmentation or detection, and subsequent tracking are the methods of choice [22]. As a result, detailed knowledge of the mechanics of morphogenetic processes can be gained [7]. In case the recorded tissue lacks structure, particle image velocimetry (PIV) is generally used to compute (sparse) displacement fields in image sequences. See, for example, [37, 47].

One difficulty in abovementioned approaches is that the observed structures often have a short life time and are being degraded during the observation, while new structures of the same type are being created [47]. Indeed, these molecular structures, and thus their fluorescent signal response, can be described by an advection–reaction equation rather than pure advection [42]. The signal variations due to reaction are a source of error in the estimation of motion that we propose to address in this paper.

Beyond the need of measurements of dense velocities of moving fluorescently-labelled molecular structures that prove robust with respect to the reaction term, it is of general interest to quantify the reaction term itself [42], and of general interest to extract information about all the physical quantities and processes, such as diffusion, that govern the observed tissue flow. For this, an accurate estimation of the velocity field corresponding to the time evolution of the distribution of fluorescent molecules is crucial [42, 47].

In this article, we argue that utilising variational motion estimation can help to identify physical quantities by estimating velocities in real data. For simplicity, we choose to demonstrate this method in a case where the biophysical data can be reduced to one space dimension. This allows to create convenient space-time representations, so-called kymographs.

One-dimensional data are indeed relevant in tissue dynamics when the cellcell junctions are found to be aligned along a straight line called a supracellular actomyosin cable [8, 40]. A common experiment to investigate the function of these cables is to cut them locally using intense laser illumination [25] and to observe the dynamics that follow. Figure 1 illustrates a prototypical two-dimensional (time-lapse) fluorescence microscopy image sequence where a cable is being severed by such a laser ablation.

**Fig. 1:**
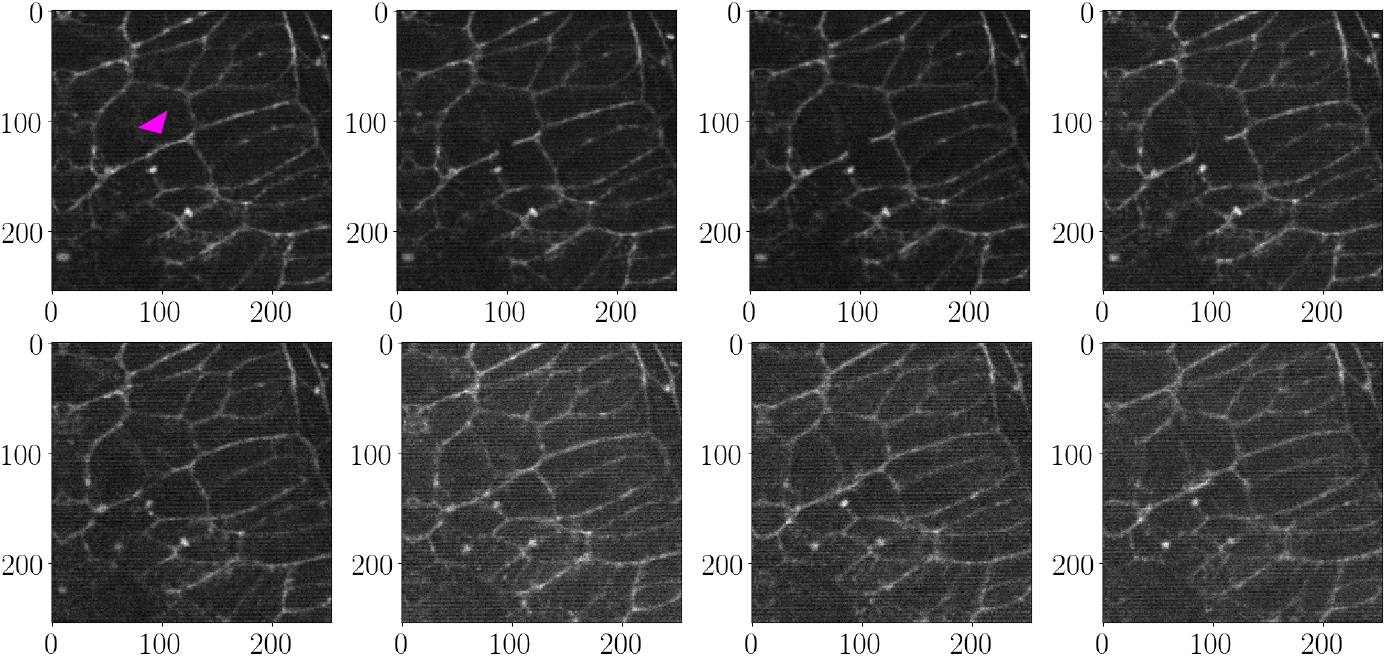
Depicted are frames no. 4, 6, 10, 20, 40, 60, 80, and 90 (left to right, top to bottom) of a 2D image sequence of fluorescently labelled cell membranes of Drosophila during a laser nanoablation. The entire sequence contains 100 frames recorded over roughly 6.5 *s*, and the imaged section spans approximately 42.2 × 42.2 μm^2^. The laser ablation is applied at frame number five, i.e. between the first and the second image shown. Its location is indicated with a magenta arrow. Observe the instantaneous recoiling and tissue loss of the cut membrane, and the subsequent growth of tissue in the cut region. Moreover, note the changes in contrast over time and the line artefacts.

Most of the relevant dynamics occur along the cable itself, thus projecting the recorded signal in a narrow stripe of less than two micrometers along its average direction preserves most of the information. See Fig. 2 for an example of a kymograph obtained from the image sequence shown in Fig. 1. Variations in fluorescence intensity are clearly visible and displacements of features can easily be measured, e.g. with existing (tracking) tools [12, 13, 38, 41]. See also Fig. 8 for an example of manually created tracks.

**Fig. 2:**
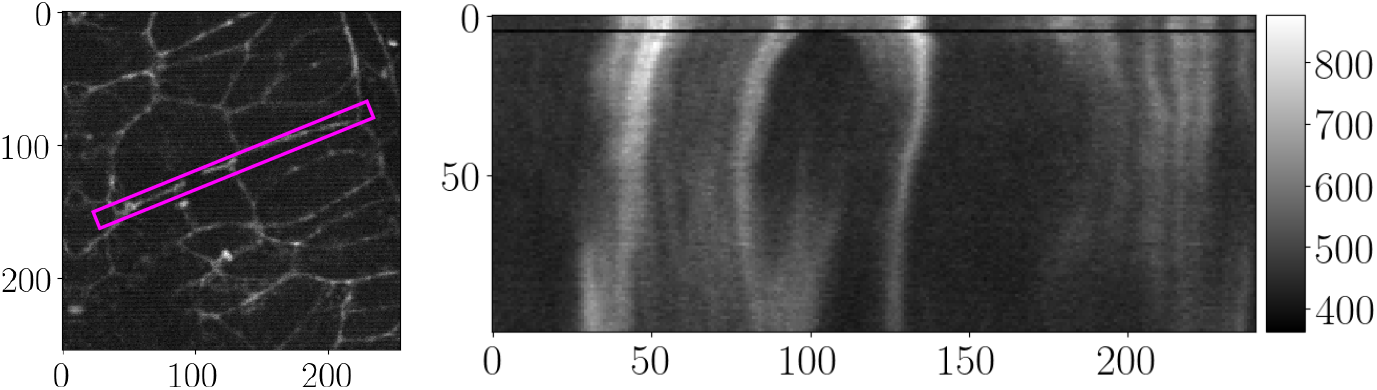
The left image shows one frame of the image sequence in Fig. 1. A supra-cellular cable is indicated with a magenta rectangle. To reduce the problem to one dimension, the data is summed along the transverse direction within the rectangular zone. The right image depicts the kymograph obtained from this dimension reduction. Time runs from top to bottom and the horizontal black line at frame five indicates the time of the laser ablation, during which the signal acquisition was paused.

Analysing such data is challenging for many reasons. First, simultaneous estimation of velocity and decay or increase of the signal renders the problem ill-posed, as we will illustrate below, and suitable (qualitative) assumptions on favoured solutions are required. Second, the obtained kymographs are very noisy, contain artefacts due to the acquisition technique, and sometimes suffer from off-plane motion of the cables. Third, the velocity field potentially contains discontinuities at the time of the laser ablation. Fourth, data is missing during the application of the laser cut and only a limited field of view is available due to the nature of the kymograph. Moreover, bleaching of tissue leads to a decrease in contrast when being exposed over long periods of time. See Figs. 1 and 2 for illustration of these issues. In this work we address mainly issues one and two.

Motivated by the laser nanoablation problem we restrict ourselves to the onedimensional case and denote by Ω ⊂ ℝ the spatial domain. For *T* > 0, we model the actomyosin concentration as a function *f*: (0,*T*) × Ω → ℝ that is proportional to the observed fluorescence response. In the following, we assume that it solves the Cauchy problem for the nonhomogeneous continuity equation in one dimension:

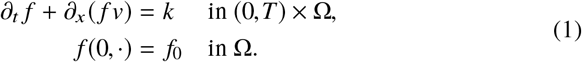

Here, *v*: (0,*T*) × Ω → ℝ is a given velocity field, *k*: (0,*T*) × Ω → ℝ a given source (or sink), and *f*_0_: Ω → ℝ a given initial condition.

Solution theory for problem (1) is closely related to solutions of the initial value problem

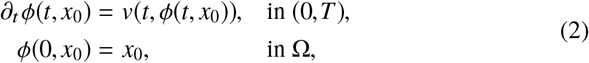

via the method of characteristics, see e.g. [24, Chap. 3.2]. Here, the map *ϕ*: [0,*T*) × Ω → Ω denotes a so-called flow of *v*. Existence and uniqueness of problem (1) can be established by application of the Picard–Lindelöf theorem to (2), provided that both *v* and *k* are sufficiently regular. In particular, it can be shown that (2) has a unique (global) solution and, moreover, that this solution is a diffeomorphism. For an introduction and for details we refer, for instance, to [18, 60].

In this work we are concerned with the solution of the inverse problem associated with (1), which is to estimate a pair (*v, k*)^⊤^ from (potentially noisy) observations *f*. Its ill-posedness is immediate because the problem is underdetermined and, given a solution (*v*_1_, *k*_1_)^⊤^, the pair (*v*_2_, *k*_1_ − *∂_x_*(*fv*_1_) + *∂_x_*(*fv*_2_))^⊤^ for *v*_2_ differentiable denotes a solution as well. In addition, it is easy to see that—without further assumptions on *v* and on *k*—the pair (0, *∂_t_ f*)^⊤^ is always a solution, albeit not a desired one.

In view of this ill-posedness and this ambiguity we consider the variational form

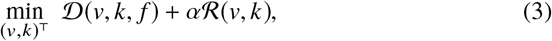

where the data term 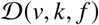 is the squared *L*^2^ norm of the first equation in (1), 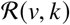 a suitable regularisation functional, and *α* > 0 is a regularisation parameter. In this article we consider different choices of 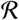. While source identification has been treated before in the literature, e.g. in [4], the main goal of this article is to recover a source (or sink) *k* which is constant along characteristics of the flow (2). In other words, we are interested in utilising the *convective derivative*

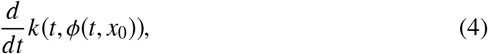

for regularisation.

Its use is inspired by the work in [33], where the convective derivative of the velocity along itself was used. From a physical perspective this choice seems natural as, in comparison to using, for instance, the *L*^2^ norm or the *H*^1^ seminorm for regularisation of *k*, it is consistent with the movement of the tracked cell tissue, which can be assumed to be the main origin of changes in the observed fluorescence intensity. However, from a numerical point of view this choice comes at the expense of having to solve nonlinear optimality conditions.

### 1.2 Contributions

The main contributions of this article are as follows. First, we propose a variational model based on the non-homogeneous continuity equation for joint motion estimation and source identification in kymographs. Second, we study the variational properties of utilising the convective derivative (4) for regularisation. Following [33], we establish existence of minimisers of the nonconvex functional. Third, for the numerical solution of the corresponding nonlinear Euler–Lagrange equations we propose to use Newton’s method and a finite element discretisation. Fourth, we present numerical results based on kymographs of laser nanoablation experiments conducted in live Drosophila embryos. Moreover, we provide an extensive experimental evaluation of different data fidelity and regularisation functionals based on manually created tracks, and evaluate our approach using synthetic data generated by solving a mechanical model of tissue formation.

### 1.3 Related Work

In [30], Horn and Schunck were the first to pursue a variational approach for dense motion estimation between a pair of images. They considered a quadratic Tikhonov-type functional that relies on conservation of brightness and used (squared) *H*^1^ Sobolev seminorm regularisation. This isotropic regularisation incorporates a preference for spatially regular vector fields. Well-posedness of the functional was proved in [51] and the problem was solved numerically with a finite element method. For a general introduction to variational optical flow see, for instance, [5].

In [57], the problem was treated on the space-time domain and extended to incorporate both spatial as well as temporal isotropic regularisation. In [56], a unifying framework for a family of convex functionals was established, and both isotropic and anisotropic variants were considered. We refer to [54] for more details on nonlinear diffusion filtering, and to [55] for an overview of numerous optical flow models and a taxonomy of isotropic and anisotropic regularisation functionals.

The convective derivative has already been used in several works. For instance, in [14] for simultaneous image inpainting and motion estimation. In [43], an optical flow term was incorporated in a Mumford–Shah-type functional for joint image denoising and edge detection in image sequences. Moreover, in [11] it was used for joint motion estimation and image reconstruction in a more general inverse problems setting. In [33] the convective acceleration was used for regularisation together with a contrast invariant Horn–Schunck-type functional. The corresponding nonlinear Euler–Lagrange equations were solved using a finite element method and alternating minimisation.

According to [16], the article [52] is credited for being the first to propose the use of the less restrictive continuity equation for motion estimation. Later it was used, for instance, to find 3D deformations in medical images [53, 20], to analyse meteorological satellite images [6, 16, 61], and to estimate fluid [15, 59] and blood flow [3] in image sequences. For a general survey on variational methods for fluid flow estimation see [29].

Whenever mass conservation is not satisfied exactly, e.g. due to illumination changes, it can be beneficial to account for these violations. For instance, in [27] they incorporated physical models. In [4] they simultaneously estimated image intensity, flux, and a potential source. In contrast to our work, only *L*^2^ integrability of the source was assumed and the constraint was enforced exactly, leading to an optimal control formulation, which was solved with a finite element method.

## 2 Problem Formulation

### 2.1 Preliminaries

#### Notation

For *T* > 0 and Ω ⊂ ℝ a bounded, connected, and open set we denote by *E* = (0,*T*)×Ω the spatio-temporal domain and by *∂E* its boundary. For a smooth function *f* : *E* → ℝ we denote by *∂_t_ f*, respectively, *∂_x_ f* the partial derivatives with respect to time and with respect to space, and by ∇*f* = (*∂_t_f, ∂_x_f*)^⊤^ its spatio-temporal gradient. The space-time Laplacian of *f* is denoted by Δ*f*. Analogously, for a smooth vector field *w*: *E* → ℝ^2^ with *w* = (*w*^1^, *w*^2^)^⊤^, its gradient is denoted by ∇*w* = (∇*w*^1^, ∇*w*^2^)^⊤^ and its spatio-temporal divergence is given by ∇ · *w* = *∂_t_ w*^1^ + *∂_x_ w*^2^. Moreover, we will write *A* ≲ *B* whenever there exists a constant *c* > 0 such that *A* ≤ *cB* holds. Finally, by the Cauchy–Schwarz inequality and application of Young’s inequality, we have

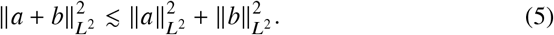

Here, and in the following, we use ║·║_*L*^2^_ instead of ║·║_*L*^2^(*E*,ℝ)_ and ║·║_*L*^2^(*E*,ℝ^2^)_ for simplicity.

#### Convective Derivative

Let *ϕ*: *E* → Ω be a flow through the domain Ω, i.e. for every *t* ∈ (0,*T*) the map *ϕ*(*t*, ·): Ω → Ω is a diffeomorphism and, for a fixed starting point *x*_0_ ∈ Ω, the trajectory *ϕ*(·, *x*_0_) is smooth.

With every trajectory *ϕ*(·, *x*_0_) that originates at *x*_0_ ∈ Ω we can associate a velocity at every time *t* ∈ (0,*T*) via (2). Thus, a flow *ϕ* gives rise to a scalar (velocity) field *v*: *E* → ℝ by means of

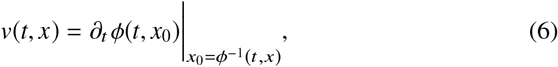

where *ϕ*^−1^(*t, x*) denotes the inverse of *ϕ*(*t*, ·) at *x* ∈ Ω, which is the starting point of the curve that passes through *x* at time *t*.

For a scalar quantity *k* : *E* → ℝ, we define the *convective derivative* of *k* along a flow *ϕ*, denoted by *D_v_k*, as

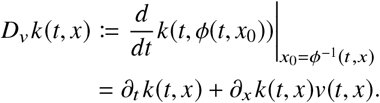

For convenience we adopt the notation 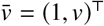. Clearly, 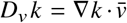 vanishes for pairs (*v, k*)^⊤^ such that 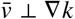 in ℝ^2^. Moreover, from a straightforward calculation we obtain

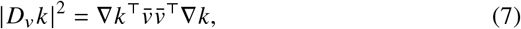

where 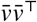 is the matrix

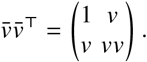

As noted in [33], the action of *D_v_k* becomes more clear by writing (7) as

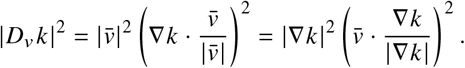

The first identity states that, for fixed 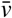, minimisation of the functional

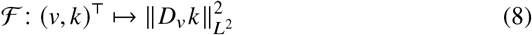

promotes functions *k* that vary little in the direction of 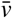. In addition, the weighting factor gives importance to regions of large 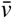. On the other hand, the second identity states that, for fixed *k*, minimisation of (8) favours functions 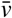 that are tangent to the level lines of *k*. This time, importance is given to regions where ∇*k* is large. See Fig. 3 for illustration.

**Fig. 3:**
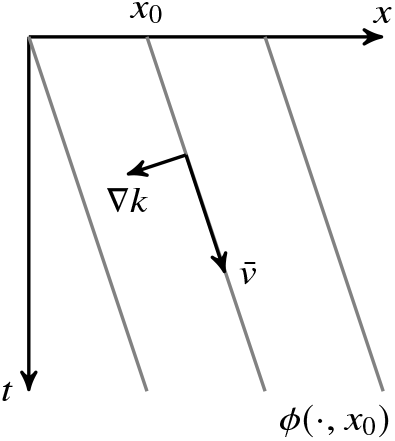
Illustration of a pair (*v, k*)^⊤^ minimising (8) where the velocity *v* is constant.

The connection to anisotropic diffusion [54] is immediate when considering the Euler–Lagrange equations associated with (8). They read

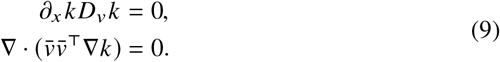

Note that the system (9) is nonlinear.

#### 2.2 Variational Model and Existence of a Minimiser

In this section we formulate the joint motion estimation and source identification problem and study the use of (8) as regularisation functional. For simplicity, we will denote a velocity-source pair by *w* = (*v, k*)^⊤^. In the following, we intend to minimise a variational formulation of the form

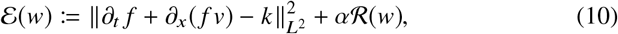

where 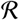 is yet to be defined and *α* > 0 is a regularisation parameter.

In order to fix a concrete choice of 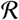 let us discuss two issues. First, it should be selected such that the functional is well-defined for an appropriate function space. In particular, we require the weak derivative of *v* in the data term in (10) to exist and to be bounded with respect to an appropriate norm. Second, *∂_x_*(*f v*) and *k* need to differ qualitatively in order to obtain a meaningful decomposition of the signal *∂_t_ f*.

Before we state the model, let us follow the ideas in [33, Chap. 5.2.2] and discuss some issues arising with the choice

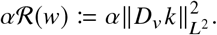

To ensure well-definedness of the functional (10) we derive, by application of (5) and HÃ¶lder’s inequality, the estimates

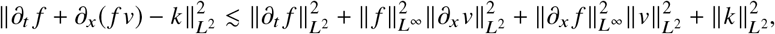

and

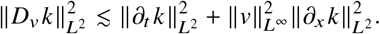

As a consequence, minimisation over the space

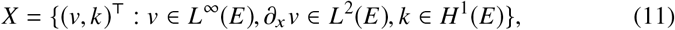

seems appropriate for data *f* ∈ *W*^1,∞^(*E*) in the space of functions with essentially bounded weak derivatives up to first order.

However, in order to show existence of minimisers of (10) by application of the direct method [19], one requires a coercivity estimate of the form

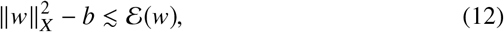

where *b* ≥ 0 is some constant and the norm of the space *X* is defined via

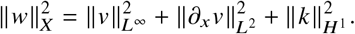

Without further restriction of the data *f*, the above functional is not coercive with respect to this space as the following example shows.

##### Example

Let *f* = *ax* for some 0 < *a* < +∞ and, consequentially, we have that *∂_x_f* = *a*. Let {*w_n_*} be the sequence with *w_n_* = (*n, an*)^⊤^, for *n* ∈ ℕ, and observe that ║*w_n_*║_*X*_ → ∞ as *n* → +∞, while the value of the functional 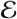 stays bounded, in fact, zero.

Moreover, it is clear that, for pairs *w* = (0,*k*)^⊤^, the inequality (12) cannot hold in general. As a remedy, we consider minimising over all pairs *w* = (*v, k*)^⊤^ arising from the Sobolev space *H*^1^(*E*, ℝ^2^). Its norm is defined via

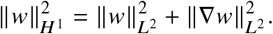

In further consequence, our goal is to find a minimiser *w* ∈ *H*^1^(*E*, ℝ^2^) of the functional 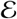 in (10) with

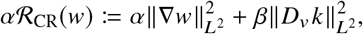

and regularisation parameters *α, β* > 0. Our final model thus reads

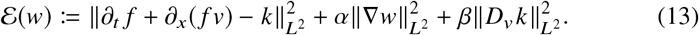

Let us establish the existence of a minimiser to the problem 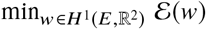. The following lemma will be used to show coercivity of 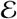 and is along the lines of [51, p. 29].

##### Lemma 1

*Let w* ∈ *L*^2^(*E*,ℝ^2^) *be constant. Then, for* ∇_−1_*f*:= (*∂_x_f*,−1)^⊤^ *with ∂_x_f* ≢ const. *the inequality*

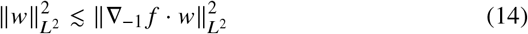

*holds*.

*Proof* First, observe that, by the Cauchy–Schwarz inequality, the assumption *∂_x_f* ≢ const. is equivalent to

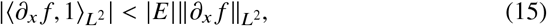

where |*E*| denotes the measure of *E*.

Next, suppose to the contrary that there is no *C* > 0 such that (14) holds. Then, for all *n* ∈ ℕ there exists *w_n_* ∈ *L*^2^(*E*, ℝ^2^) such that

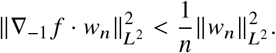

Let 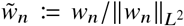 with 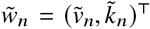 and obtain 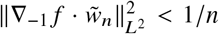. But then,

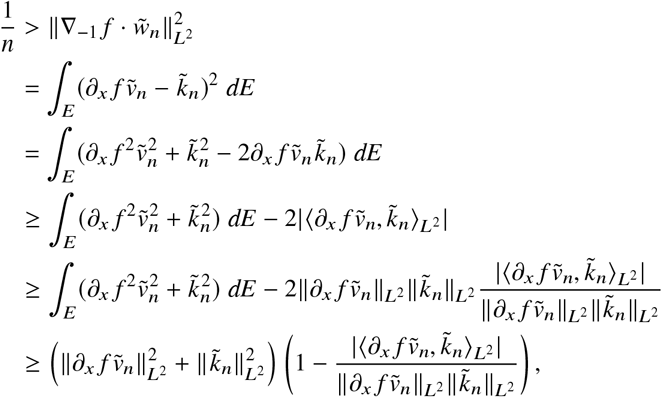

together with (15) implies that 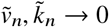 as *n* → +∞, contradicting the assumption. Therefore, (14) holds.

In particular, Lemma 1 holds for the (componentwise) average *w_E_* of *w*, defined as

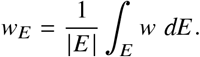

The following proposition is a straightforward adaptation of [33, Prop. 1] and utilises the direct method in the calculus of variations [19].

##### Proposition 1

*For f* ∈ *W*^1,∞^(*E*) *satisfying* (15), *the functional 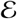 admits a minimiser in H*^1^(*E*, ℝ^2^).

*Proof* The functional is proper and bounded from below since all terms are nonnegative and, for w identically zero,

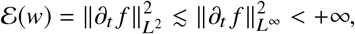

since *f* ∈ *W*^1,∞^(*E*).

Next, we show coercivity of (13) with respect to *H*^1^(*E*, ℝ^2^). Observe that

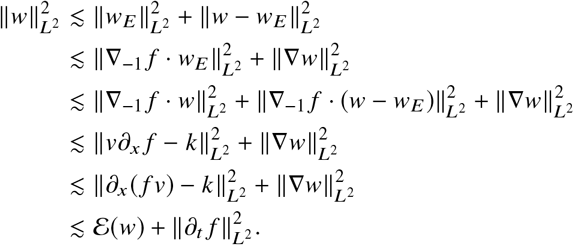

The chain of inequalities follows from (5), (14), the Poincaré–Wirtinger inequality [24, Chap. 5.8],

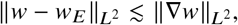

and the assumption *f* ∈ *W*^1,∞^(*E*). Coercivity of 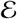 with respect to *H*^1^(*E*,ℝ^2^) then follows since 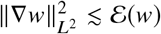.

Next, we discuss sequential weak lower-semicontinuity of 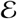. Let {*w_n_*} ⊂ *H*^1^(*E*, ℝ^2^) such that *w_n_* ⇀ *ŵ* in *H*^1^(*E*, ℝ^2^). In particular, we have that ∇*w_n_* ⇀ ∇*ŵ* in *L*^2^(*E*,ℝ^4^). For 1 < *p* < 2, the compact embedding *H*^1^(*E*,ℝ^2^) ⊂ *W*^1,*p*^(*E*,ℝ^2^) ⊂⊂ *L*^2^(*E*,ℝ^2^) holds, see [24, Chap. 5.7]. As a consequence, there exists a subsequence, also denoted by *w_n_*, such that *w_n_* → *ŵ* in *L*^2^(*E*, ℝ^2^). Then, weak lower-semicontinuity of 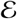 follows by application of [19, Thm. 3.23] since, for fixed *w*, we have that all terms are convex in ∇*w*. In particular, |*D_v_k*|^2^ is a quadratic form and therefore convex in ∇*w*, since 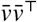 in (7) is symmetric positive semidefinite.

Finally, by application of [19, Thm. 3.30], the functional 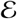 admits a minimiser.

Let us add, however, that the convective regularisation functional 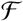 is nonconvex, as the following example shows. We pick *w*_1_ = (0, *x*)^⊤^ and *w*_2_ = (1, 0)^⊤^, and obtain

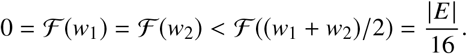

Therefore, 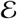 is nonconvex in general and several minima might exist. However, for *β* = 0 a unique minimiser exists.

## 3 Numerical Solution

In this section we derive necessary conditions for minimisers of (13) and discuss the numerical solution of a weak formulation by means of Newton’s method.

### 3.1 Euler–Lagrange Equations

For convenience let us abbreviate *F* := *∂_t_f* + *∂_x_*(*fv*) − *k*. The Euler–Lagrange equations [17, Chap. IV] associated with minimisation of the functional 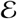 in (13) then read

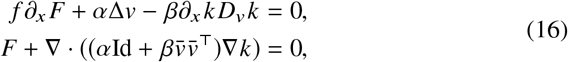

where Id denotes the identity matrix of size two. Recall from Sec. 2.1 that Δ and ∇ are spatio-temporal operators. Moreover, the natural boundary conditions at *∂E* are given by

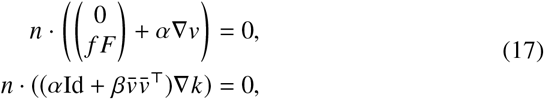

where *n* ∈ ℝ^2^ is the outward unit normal to the space-time domain *E*.

Let us highlight two aspects of (16). First, note that the system is nonlinear in the unknown *w* = (*v, k*)^⊤^ due to the convective regularisation. Second, as already mentioned in Sec. 2.1, there is a connection of the second set of equations in (16) with anisotropic diffusion with the diffusion tensor given by 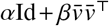. The investigation of existence and regularity of solutions of (16) is left for future research.

### 3.2 Weak Formulation and Newton’s Method

We minimise 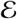 by applying Newton’s method to the weak formulation associated with (16) together with boundary conditions (17). It is derived as follows.

Multiplying with a test function *φ* = (*φ*^1^, *φ*^2^)^⊤^ ∈ *H*^1^(*E*, ℝ^2^), integrating by parts under consideration of (17), and adding both equations leads to the following variational problem: Find *w* ∈ *H*^1^(*E*, ℝ^2^) such that

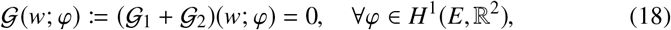

with

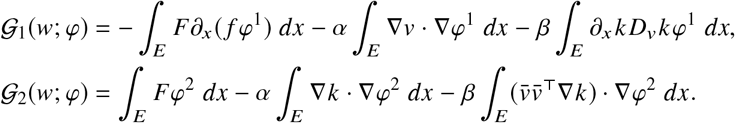

The Gâteaux derivative 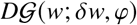 of 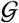 at *w* ∈ *H*^1^(*E*, ℝ^2^) in the direction of *δw* = (*δv, δk*)^⊤^ ∈ *H*^1^(*E*, ℝ^2^) is given by 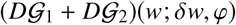 with

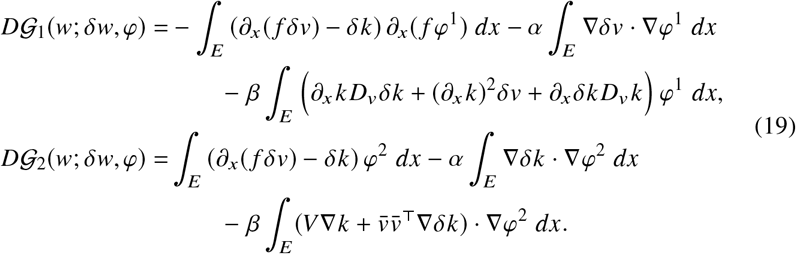

Here, *V* is the Gâteaux derivative of 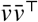 at *v* in the direction *δv* and reads

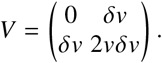

We solve the nonlinear problem (18) with Newton’s method, which proceeds as follows. Starting from an initial solution *w*^(0)^ = (0, 0)^⊤^ we update the solution according to the rule

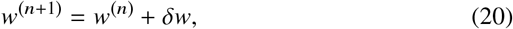

where *δw* is the update and *n* ∈ ℕ_0_. Computing *δw* in each step requires to solve a linear variational problem: Find *δw* ∈ *H*^1^(*E*, ℝ^2^) such that

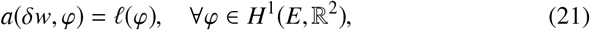

with

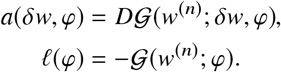

Note that the dependence on the current iterate *w*^(*n*)^ is through 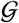 and 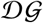, defined in (18) and (19), respectively. In our experiments we observed that (20) typically converges within only a few iterations.

### 3.3 Discretisation

We have implemented the weak formulation in FEniCS [1], which can also handle Newton’s method automatically. The formulation (18) was discretised using multilinear finite elements. Since kymographs serve as input data we have discretised the rectangular space-time domain *E* with a triangular mesh based on the regular grid so that every vertex of the mesh corresponds to one pixel of the image *f* and to one pair of values of the unknown *w*.

Moreover, in the implementation we penalise weak derivatives differently in space and time. This results in four regularisation parameters 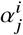, with *i* = {*v, k*} and *j* = {*t, x*}, for the *H*^1^ seminorm in (13) and one additional parameter *β* for the convective regularisation. Due to the equivalence of norms the existence result in Prop. 1 still holds true.

Integrals are computed exactly with an appropriate Gauss quadrature, which is automatically selected by FEniCS. Similarly, since the image *f* is represented by a piecewise multilinear function, products of partial derivatives of *f* that appear on the right-hand side of (21) are automatically projected onto the correct space.

As termination criterion for Newton’s method we used the default criteria of FEniCS with both the absolute and the relative residual of (18) set to 10^−10^. The maximum number of iterations was set to 15. Convergence was typically achieved within only a few iterations, which usually amounted to just a few seconds of computing time on a standard consumer laptop. It needs to be mentioned that, in our experiments we found that, the method fails to converge when the parameters 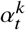 and 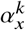 are chosen too small in comparison to *β*. This is, however, in line with the theoretical results in Sec. 2.2, which require *H*^1^ regularity of *k*.

Both the source code^1^ of our Python implementation and the microscopy data^2^ used in the experiments are available online.

## 4 Experimental Results

In this section we present numerical results. In the first part, we demonstrate the importance of estimating a source based on synthetic data. Then, in the second part we show results for nonsynthetic microscopy data. After briefly discussing the data, we investigate the effects of varying the regularisation parameter *β*, which controls the amount of convective regularisation. We then qualitatively compare the (standard) 1D variational optical flow model with our continuity equation-based formulation for several choices of the regularisation functional 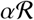 in (10). In addition, we present a quantitative evaluation of the considered models based on recoil velocities obtained from manual tracking of features in kymographs. Finally, in the last part, we evaluate our approach based on data coming from the solution of a mechanical model of tissue formation.

In all results, the computed velocity fields are presented visually with the help of streamlines, see e.g. [58]. These are integral curves computed by numerically solving the ordinary differential equation (2) for the estimated velocity *v* and a selected number of initial points. The resulting curves are then colour coded according to their velocities and shown superimposed with the kymograph data. This representation is more comprehensible and allows to visually check whether the estimated velocities are approximately correct.

### 4.1 Analytical example

In order to demonstrate the necessity of estimating a source when the changes in the signal cannot be explained using mass conservation we conducted experiments for synthetic data. To this end, we generated a signal *f*, given as

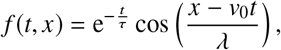

on a periodic domain Ω = (0, 1) and for the time interval [0, 1]. The parameters were set to *τ* = 1 and to *λ* = 1/(4*π*). This signal shifts to the right with constant velocity *v*_0_ = 0.1 and decays exponentially over time in its magnitude. It can easily be verified that the source is given by *k*(*t, x*) = −*f*(*t, x*)/*τ*.

We then solved the variational problem using in one case the homogenous and in one case the non-homogenous continuity equation. Periodic boundary conditions in space were enforced, and regularisation parameters were set to 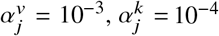, for *j* ∈ {*t, x*}, and to *β* = 10^−3^. Since *k* is not constant along characteristics in this example we haven chosen *β* quite small.

Figure 4 illustrates the results of this experiment. It can clearly be seen that not accounting for the decay of the signal leads to a velocity that is significantly different to v0 in this example, whereas using the non-homogenous continuity equation estimates both the velocity and the source (not shown) very well.

**Fig. 4:**
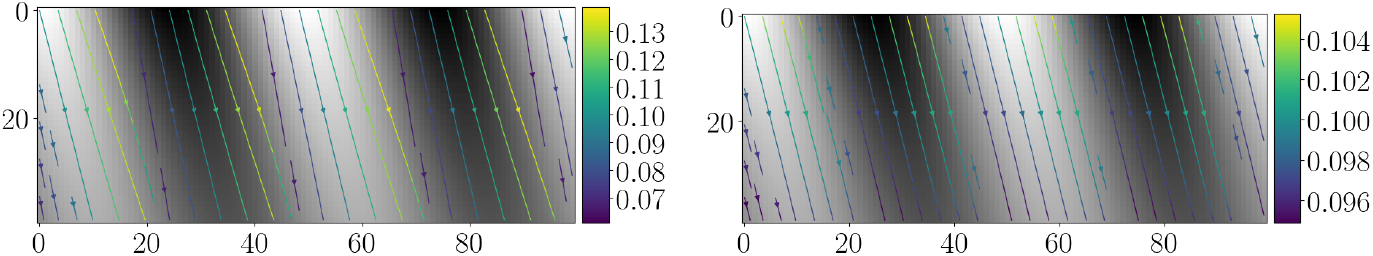
In this example, we demonstrate the necessity of source estimation for decaying data. Here, we have generated a signal that shifts to the right with constant velocity *v*_0_ = 0.1 and simultaneously decays exponentially (in magnitude). Shown are the signal *f* and streamlines computed from the estimated velocities using the homogenous (left) and the non-homogenous (right) continuity equation.

### 4.2 Microscopy Data and Acquisition of Kymographs

The data at hand are 2D image sequences of living Drosophila embryos recorded with confocal laser-scanning microscopy. We refer to [48] for the used microscopy technique and for the preparation of flies, as well as for the details of the laser ablation method. For this study we recorded 15 image sequences, all of which feature cell membranes that have been fluorescently labelled with E-cadherin:GFP, see [48]. The image sequences feature a square region of approximately 42.2 × 42.2 μm^2^ at a spatial resolution of 250 × 250 pixels. A typical sequence contains between 60 and 100 frames that were recorded at a temporal interval of 727.67 ms, and the recorded image intensities *f*^2D+T^ are in the range {0,…, 255}.

Each of the sequence shows a single plasma-induced laser nanoablation, which led to the controlled destruction of tissue in a linear region of 2 μm length that is approximately orthogonal to the cable. This ablation is expected to have a width of the order of the size of one pixel. Recall that in Fig. 1 we show such a typical dataset.

In order to obtain a kymograph from each microscopy sequence, we first labelled the location of the intersection between the ablation line and the actomyosin cable with a point *c* ∈ ℝ^2^. Then, we visually determined an approximate orientation, given by a unit vector *e* ∈ ℝ^2^, of the selected cable by defining a straight line of length 2*L* + 1 pixels which passes through *c*. Typically, *L* = 100 pixels is sufficient for the considered datasets.

To create a one-dimensional image sequence we used standard nearest neighbour sampling along the abovementioned line. A nearest neighbour in terms of the pixel locations *P* of *f*^2D+T^ is given by

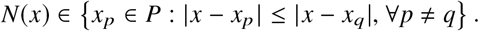

Then, the sampling points that are separated by distance one and lie along the abovementioned straight line are given by *p_i_* = *c* + *ie*, for *i* = −*L*,…, *L*. A nearest neighbour interpolation of *f*^2D+T^ along this line is given by *f*^2D+T^(*t, N*(*p_i_*)).

However, due to noise, the spatial extent, and minor displacements of the cable in orthogonal direction, we also considered sampling points that lie on the parallel line of distance *j* = −*h*,…, *h*. These points are given by

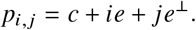

Then, at *i* = 1,…, 2*L* + 1, we define the intensity of a kymograph *f^δ^*(*t, i*) as

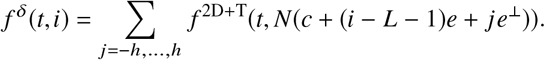

In other words, the intensity at *i* is given as the sum of nearest neighbour interpolations at points that lie on an orthogonal straight line. The superscript *δ* indicates noise in the created kymograph. We found that *h* = 5 pixels generates satisfactory kymographs.

For this process, we used the reslice tool in Fiji [49] with a slice count of 2*h* and no interpolation selected, after manually placing a straight segment along a selected actomyosin cable, see Fig. 2 for illustration. Subsequently, a projection with the `SUM` option selected creates the final kymograph.

Since the image acquisition is paused during the laser ablation, see Fig. 2, we simply replaced the missing frame with the previous one. Moreover, we applied a Gaussian filter to the kymograph *f^δ^* to guarantee the requirements specified in Sec. 2.2 and scaled the image intensities to the interval [0, 1]. The kernel size of the Gaussian filter was chosen as 10 × 10 pixels and the standard deviation set to *σ* = 1. The filtered and normalised kymograph is denoted in the following by *f*.

### 4.3 Qualitative Comparison

In the first experiment, we investigated the effect of the convective regularisation for one chosen kymograph. To this end, we solved the necessary conditions (16) as outlined in Sec. 3 for varying regularisation parameter *β* Figure 5 shows streamlines for the estimated velocities together with the computed sources. Since we did not find any significant difference between the results for *β* = 10^−4^ and for *β* = 0, we simply omit the latter.

**Fig. 5:**
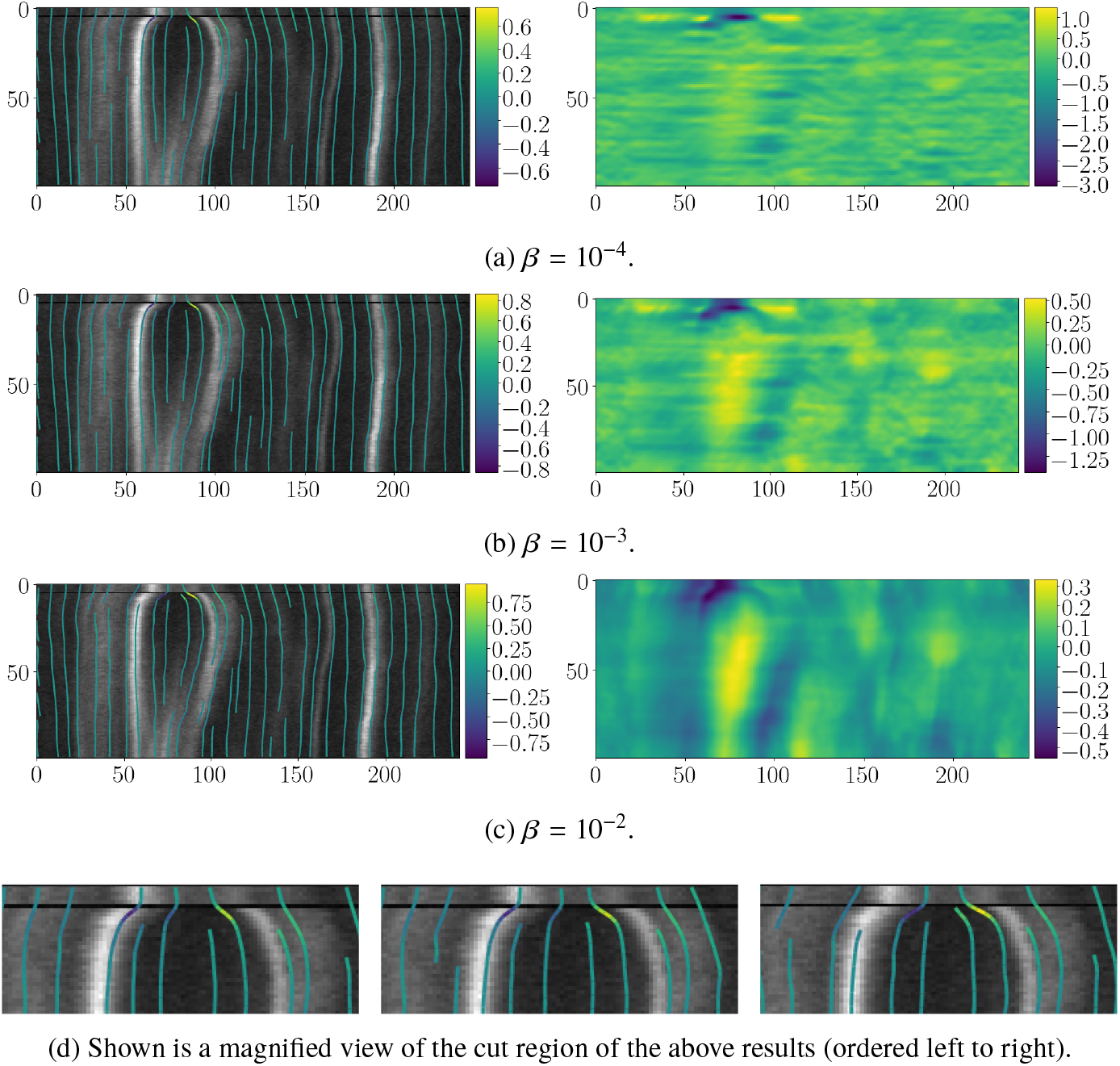
Visualisation of the effect of the convective regularisation. Shown in 5a–5c are kymographs *f^δ^* (left) with streamlines superimposed and the estimated source *k* (right) for increasing regularisation parameter *β* (from top to bottom). The other parameters were fixed and set to 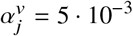 and 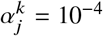, for *j* ∈ {*t, x*}.

The two main findings of this experiment are as follows. First, for the chosen dataset, the convective regularisation can help to estimate more accurate velocities shortly after the ablation was applied. As can be seen in Fig. 5c (left) and Fig. 5d (right), this leads to a more accurate estimation of the recoil velocities at the cut ends. In Fig. 5d we display a magnified view of the results in the cut region.

Second, as expected, with increasing *β* the estimated source gets more regular in the direction of the flow, see Figs. 5a-5c (right). In particular, the oscillations in space and time in the estimated source, which can be most likely attributed to noise and to artefacts created during the acquisition, decrease significantly. This certainly can lead to better interpretability and help to get a better understanding of the estimated reaction term *k*. Observe also the decrease in the magnitude of *k* as *β* increases. This is made apparent by the colour coding.

In the next experiment, we qualitatively compare the standard variational optical flow model with *H*^1^ seminorm regularisation and the continuity equation-based model in (10) paired with different regularisation functionals. The first model we consider is based on the optical flow equation [30] in one space dimension, which reads

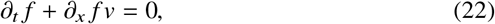

and assumes that *f* is constant along characteristics of the flow (2). Even though no source *k* is estimated in this model, the main motivation for its inclusion in the evaluation is that it can serve as a baseline method. The quantitative evaluation in Sec. 4.4 is based on manually created tracks that follow highly visible features in the kymographs and, in many cases, roughly preserve their intensity as the tissue deforms. See Fig. 8 for illustration.

We highlight that, given *∂_x_f* ≠ 0, equation (22) admits a unique solution, namely *v* = −*∂_t_f* /*∂_x_f*. However, due to noise degradation and aforementioned artefacts it is beneficial to solve (22) in a variational framework. Therefore, we consider minimising the functional

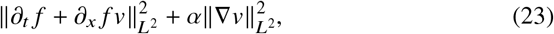

analogously to the solution method outlined in Sec. 3. In contrast to the functional (13), the corresponding Euler–Lagrange equations are linear and the weak formulation can be solved directly.

All other functionals we investigate are based on (10) and read

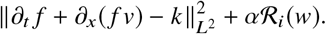

Here, 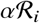 represent different choices in the regularisation and, consequentially, also in the function space we minimise over. Since the above functional doesn’t require any Sobolev regularity of the source *k*, we investigate the setting where *v* ∈ *H*^1^(*E*) and *k* ∈ *L*^2^(*E*), that is

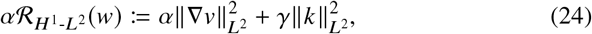

with *γ* > 0. Moreover, we also consider the choice *v* ∈ *H*^1^(*E*) and *k* ∈ *H*^1^(*E*). The regularisation functional then reads

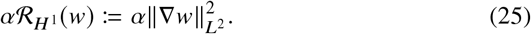

Finally, we will present results for the setting *v* ∈ *H*^1^(*E*) and *k* ∈ *H*^1^(*E*) together with the convective regularisation. This is the main model investigated in Sec. 2.2 and is given by

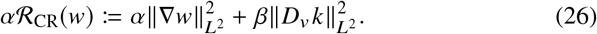

In Fig. 6 we show minimising functions for models (23)–(26) and in Fig. 7 we display a magnified view of the cut region. For all models, the regularisation parameters were chosen manually so that the streamlines obtained from the computed velocities best matched the cut ends which resulted from the laser ablation. For comparison, we also refer the reader to Fig. 8, which shows manually tracked cut ends. The four main observations from this experiment were as follows.

**Fig. 6:**
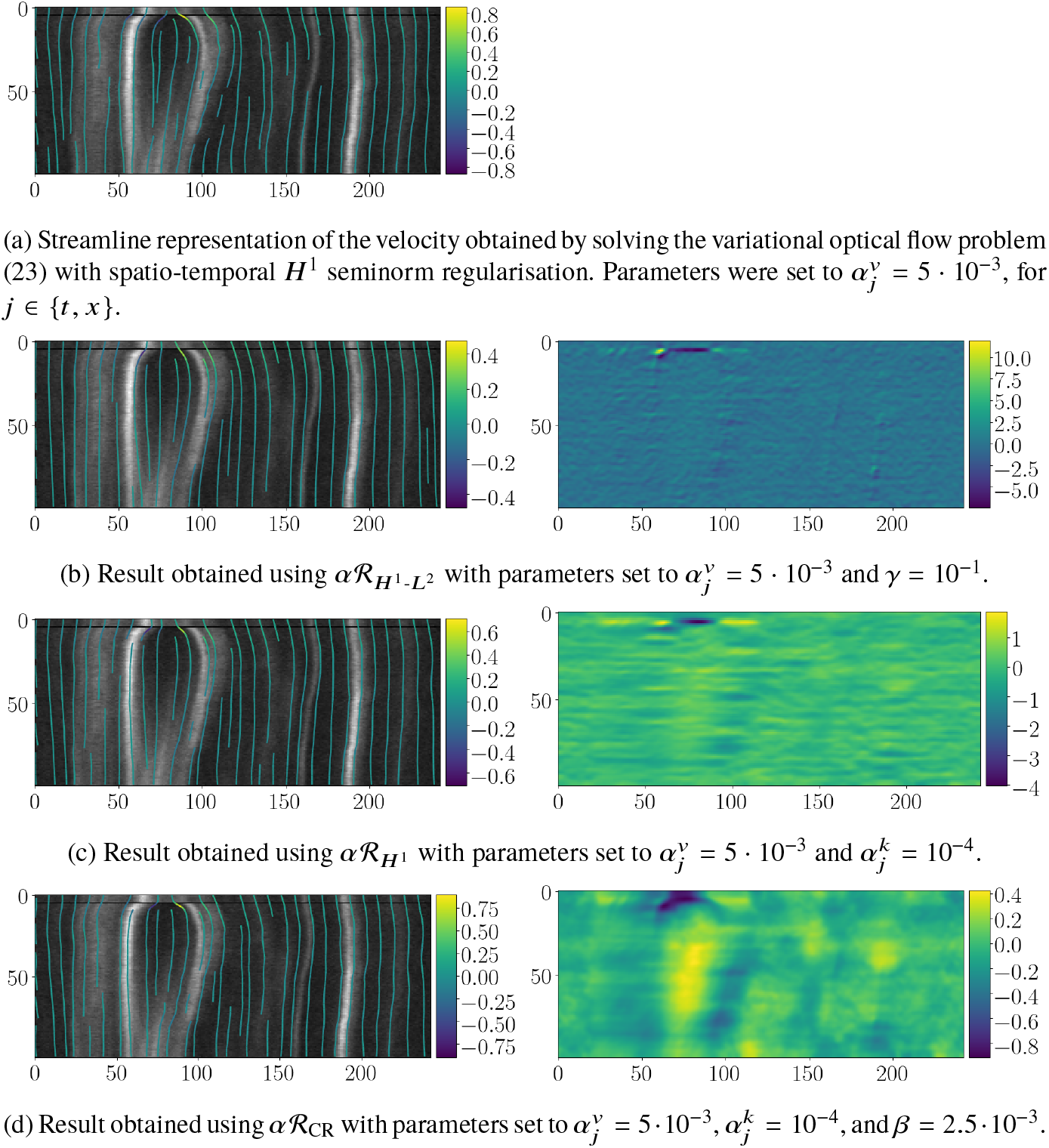
Qualitative comparison of different models based on one chosen dataset. Regularisation parameters were chosen manually so that the recovered streamlines best matched the cut ends.

**Fig. 7:**
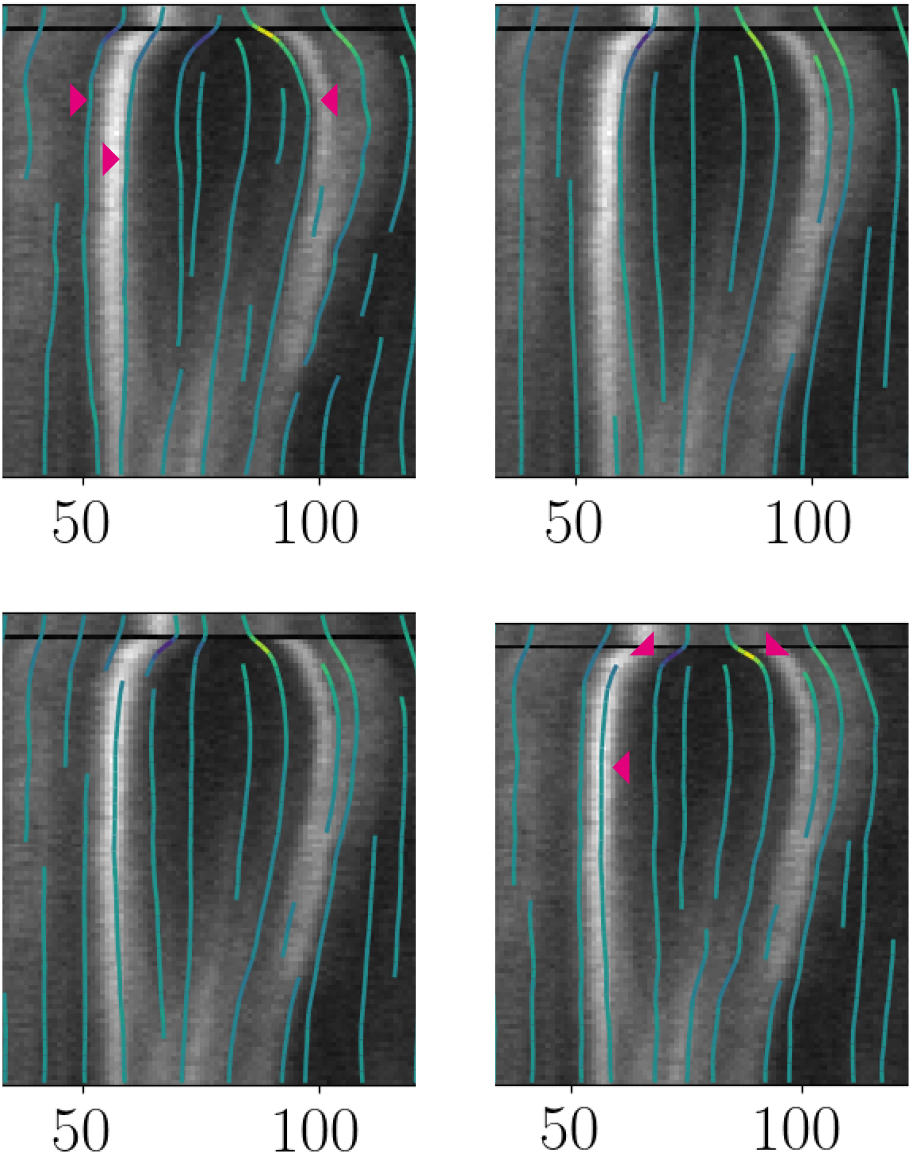
Shown is a magnified view of the cut region of the results shown in Fig. 6 (ordered left to right, top to bottom).

**Fig. 8:**
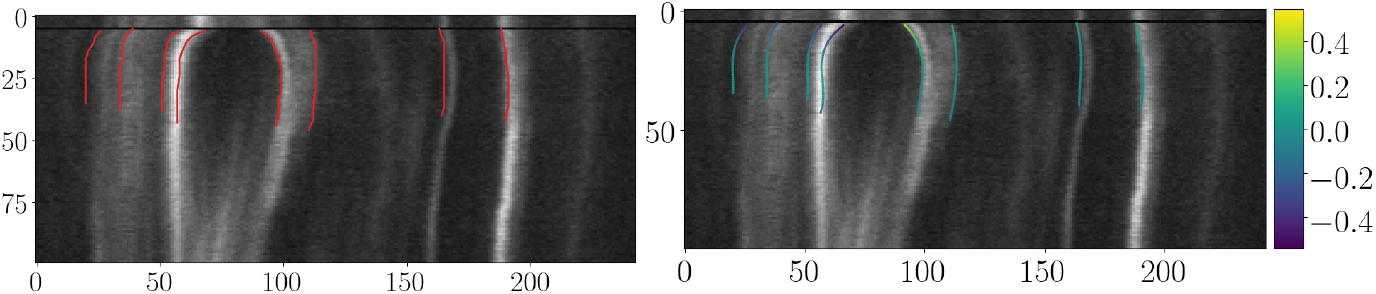
Visualisation of manually created tracks of features (left) used for the evaluation superimposed with the corresponding kymograph and their corresponding third-order spline representation (right). Colour in the spline representation indicates velocity.

First, in Fig. 6a, which shows a minimising function of the optical flow model (23), it is clearly visible that many characteristics follow paths of constant fluorescence intensity. See also the two outermost markers in Fig. 7 (left). However, this model underestimate the recoil velocity shortly after the cut and may lead to wrong characteristics, see the middle marker in Fig. 7 (left).

Second, in Fig. 6b we illustrate a minimising pair (*v, k*)^⊤^ of the continuity equation-based model using 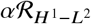. It is apparent that *k* captures both noise and artefacts, and leads to an undesirable underestimation of the velocities in the cut region. See also Fig. 7 (second from left). While this model unsurprisingly results in the smallest residual error (in norm) of the non-homogenous continuity equation, cf. also Table 1, it is insufficient for a meaningful quantification of tissue loss and growth.

**Table 1:** Residual error for every dataset and every model investigated. For each dataset we show the smallest residual in *L*^1^ norm that is obtained during the parameter search.

| Dataset | OF | $\alpha\mathcal{R}_{H^1-L^2}$ | $\alpha\mathcal{R}_{H^1}$ | $\alpha\mathcal{R}_{CR}$ |
| --- | --- | --- | --- | --- |
| 1 | 0.45 | **0.16** | 0.34 | 0.34 |
| 2 | 0.65 | **0.27** | 0.52 | 0.53 |
| 3 | 0.44 | **0.17** | 0.36 | 0.36 |
| 4 | 0.36 | **0.13** | 0.26 | 0.26 |
| 5 | 0.62 | **0.26** | 0.49 | 0.49 |
| 6 | 0.49 | **0.17** | 0.36 | 0.37 |
| 7 | 0.56 | **0.20** | 0.42 | 0.43 |
| 8 | 0.50 | **0.16** | 0.38 | 0.40 |
| 9 | 0.29 | **0.12** | 0.24 | 0.25 |
| 10 | 0.43 | **0.17** | 0.33 | 0.34 |
| 11 | 0.42 | **0.16** | 0.30 | 0.31 |
| 12 | 0.32 | **0.12** | 0.23 | 0.23 |
| 13 | 0.52 | **0.20** | 0.37 | 0.38 |
| 14 | 0.64 | **0.26** | 0.51 | 0.52 |
| 15 | 0.33 | **0.11** | 0.22 | 0.23 |
| Average | 0.47 | **0.18** | 0.36 | 0.36 |

Third, in Fig. 6c we depict a minimising pair for the model using 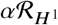 as regularisation functional. Less noise is picked up by the source *k* and the recovered velocities *v* are closer to what one would expect in the cut region, see also Fig. 7 (third image from left). However, undesired oscillatory patterns are present in the source *k*.

Finally, in Fig. 6d we show a minimiser of the model using convective regularisation, that is 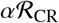. This particular choice leads to significant visual improvement both in the recovered velocity and the estimated source. Specifically, as the streamlines in the cut region right after the laser ablation in Fig. 7 show (see top markers), the movement of the tissue is captured very well. In addition, bright features such as the left cut end, cf. the bottom marker in Fig. 7, are followed accurately. Moreover, the changes in the fluorescence intensities are indicated nicely in the visualisation of the source *k*, see Fig. 6d (right). In particular, the significant increase between the cut ends towards the end of the sequence, possibly due to wound healing, is indicated adequately.

In summary, it can be said that our variational model based on the non-homogenous continuity equation together with convective regularisation can lead to improved results compared to existing models when parameters are selected by hand and comparison is performed visually.

### 4.4 Quantitative Comparison based on Measured Recoil Velocities

In this section, we compare the models (23)–(26) presented in the previous section from a quantitative point of view. Our evaluation is based on manually measured recoil velocities of clearly visible features in the kymographs, and include the cut ends resulting from the laser nanoablation.

For this comparison, we have annotated 15 kymographs in Fiji [49] to obtain discrete trajectories to compare to. See Fig. 8 (left) for an example. These tracks can then be used to compare to either the estimated velocities or to the computed characteristics that solve (2) numerically. Observe in Fig. 8 that all trajectories start only after the ablation, which is the main region of interest from a tissue mechanics point of view.

Before presenting the comparison, let us briefly discuss the methodology and the used evaluation criteria. Since some created tracks do not feature a coordinate for each time instant we interpolated each track *i* with a third-order spline *ϕi*. As a result, velocities *∂_t_ ϕ_i_* can be computed conveniently and used for comparison to the velocities estimated by the variational approach. See Fig. 8 for an example of tracks (left) and their corresponding spline interpolations (right), which are colour coded according to their velocities.

In our experiments we found that each kymograph requires the regularisation parameters to be adjusted individually. Therefore, in the experimental comparison, we performed a search over all parameter combinations

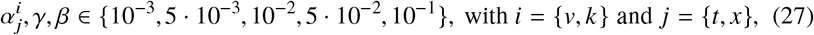

for whichever parameters are applicable to the respective model. In the case of the most complex model, i.e. the one that includes the convective regularisation (26), this amounted to probing 5^5^ parameter combinations per dataset. For each of the criteria listed below, we recorded the best result that was obtained with each model and for each kymograph during the parameter search.

#### Error in Residual

In the first comparison, the goal was to see which model best fits the recorded data. In Table 1 we report the *L*^1^ norm of the smallest observed residual of the underlying model equation, that is

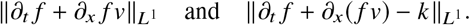

As expected, the model 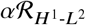 results on average and for every single dataset in the smallest residual. However, as already mentioned in Sec. 4.3, due to the high noise level it is only of limited use for quantifying the reaction term as *k* captures a significant amount of noise. The main finding of this experiment is that the optical flow model is by far not capturing the entire essence of the dataset, which is indicated by the high residual in comparison to the continuity equation-based models. Moreover, let us also highlight that the residual is on average not significantly increased in comparison to the 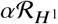 model when the convective regularisation is used in addition.

#### Error in Velocity

In the second comparison, we looked at the absolute error between the velocity of each manually created track and the velocities estimated with our models. For a particular track *ϕ_i_* of a dataset, we define this error at time *t* ∈ [0,*T_i_*] as

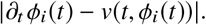

Here, *T_i_* > 0 is the length of the track and we have assumed for simplicity that all trajectories start at *t* = 0. The velocity *v* needs to be interpolated, since *ϕ_i_* is a spline representation.

In further consequence we computed, for each dataset and for each parameter configuration, the mean squared *L*^2^ norm of the error in velocity along all its *N* tracks. It is given by

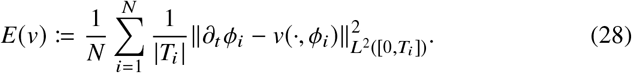

In our experiments, in none of the tested models we could find a single parameter combination that worked well for most datasets in terms of the mean error *E*(*v*). This can be seen, for example, in Fig. 9, where we have plotted exemplary for each dataset and for each parameter setting in (27) the error *E*(*v*) for the model 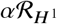.

**Fig. 9:**
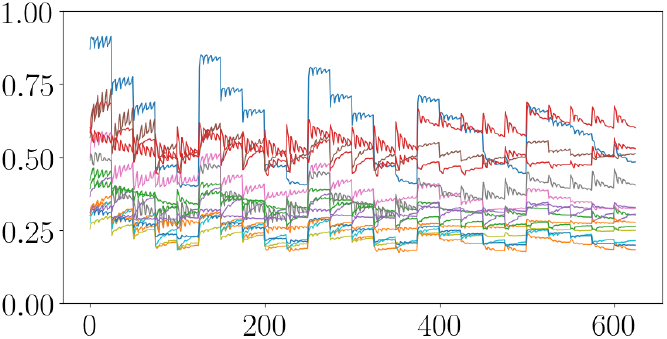
Plotted is the mean error (28) (vertical axis) for the model 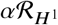 for each of the 15 datasets (in different colours) and all probed parameter combinations (horizontal axis). It can be seen that no single tested parameter settings led to a small error for all datasets.

As a consequence, we report in Table 2 (left) the best mean error *E*(*v*) that we obtained for each dataset by the grid search. The main findings are as follows. First, and most importantly, the continuity equation-based model with convective regularisation performed on average as well as the optical flow-based model when using *E*(*v*) as evaluation criterion, with the advantage of simultaneously yielding an estimate of the source.

**Table 2:**
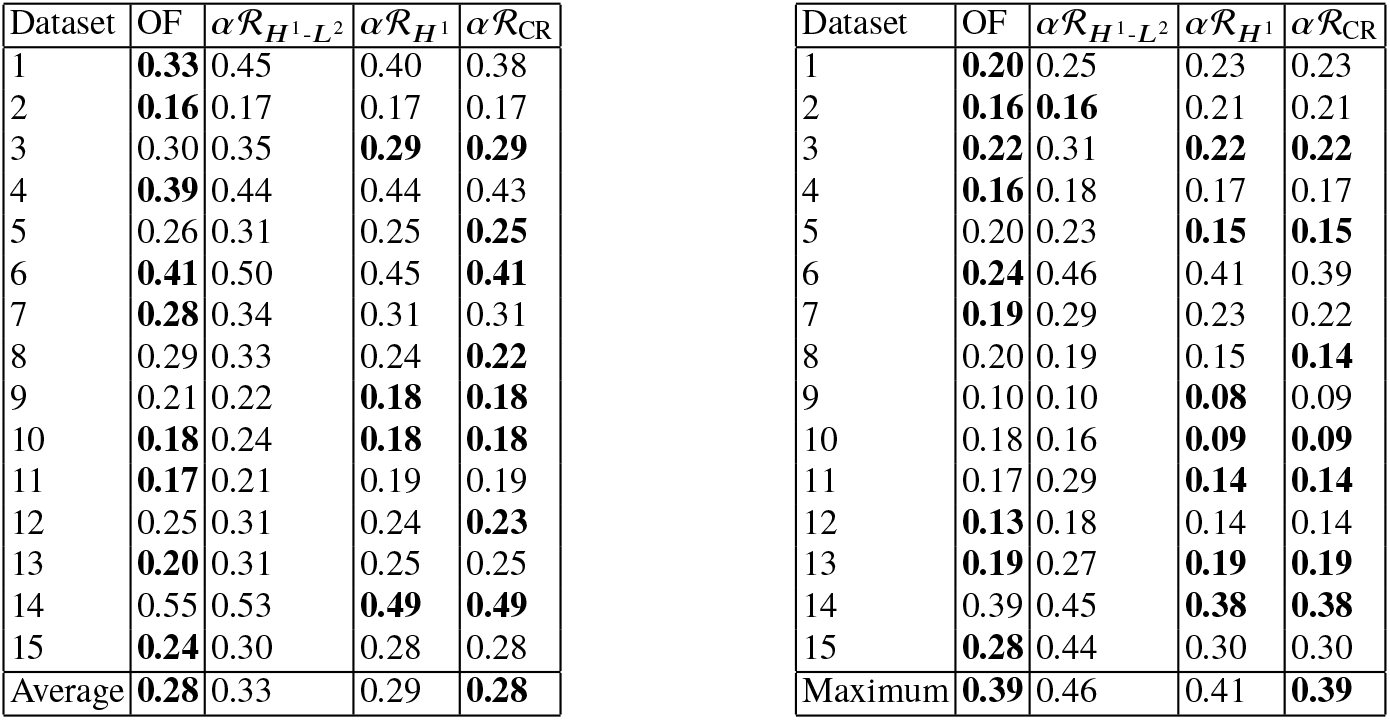
Tables depict the average error *E*(*v*) in the *L*^2^ norm (left), as in (28), and the maximum error (right) for every dataset and every model investigated. For each dataset we show the smallest error that is obtained during the parameter search.

| Dataset | OF | $\alpha\mathcal{R}_{H^1-L^2}$ | $\alpha\mathcal{R}_{H^1}$ | $\alpha\mathcal{R}_{CR}$ |
| --- | --- | --- | --- | --- |
| 1 | **0.33** | 0.45 | 0.40 | 0.38 |
| 2 | **0.16** | 0.17 | 0.17 | 0.17 |
| 3 | 0.30 | 0.35 | **0.29** | **0.29** |
| 4 | **0.39** | 0.44 | 0.44 | 0.43 |
| 5 | 0.26 | 0.31 | 0.25 | **0.25** |
| 6 | **0.41** | 0.50 | 0.45 | **0.41** |
| 7 | **0.28** | 0.34 | 0.31 | 0.31 |
| 8 | 0.29 | 0.33 | 0.24 | **0.22** |
| 9 | 0.21 | 0.22 | **0.18** | **0.18** |
| 10 | **0.18** | 0.24 | **0.18** | **0.18** |
| 11 | **0.17** | 0.21 | 0.19 | 0.19 |
| 12 | 0.25 | 0.31 | 0.24 | **0.23** |
| 13 | **0.20** | 0.31 | 0.25 | 0.25 |
| 14 | 0.55 | 0.53 | **0.49** | **0.49** |
| 15 | **0.24** | 0.30 | 0.28 | 0.28 |
| Average | **0.28** | 0.33 | 0.29 | **0.28** |
| 1 | **0.20** | 0.25 | 0.23 | 0.23 |
| 2 | **0.16** | **0.16** | 0.21 | 0.21 |
| 3 | **0.22** | 0.31 | **0.22** | **0.22** |
| 4 | **0.16** | 0.18 | 0.17 | 0.17 |
| 5 | 0.20 | 0.23 | **0.15** | **0.15** |
| 6 | **0.24** | 0.46 | 0.41 | 0.39 |
| 7 | **0.19** | 0.29 | 0.23 | 0.22 |
| 8 | 0.20 | 0.19 | 0.15 | **0.14** |
| 9 | 0.10 | 0.10 | **0.08** | 0.09 |
| 10 | 0.18 | 0.16 | **0.09** | **0.09** |
| 11 | 0.17 | 0.29 | **0.14** | **0.14** |
| 12 | **0.13** | 0.18 | 0.14 | 0.14 |
| 13 | **0.19** | 0.27 | **0.19** | **0.19** |
| 14 | 0.39 | 0.45 | **0.38** | **0.38** |
| 15 | **0.28** | 0.44 | 0.30 | 0.30 |
| Maximum | **0.39** | 0.46 | 0.41 | **0.39** |

A possible explanation for the comparably good performance of the optical flow-based model is that the manually created tracks approximately constitute trajectories of constant intensities rather than the characteristics associated with (1). However, in combination with the findings presented in Table 1, which show that the average residual is much smaller when using a continuity equation-based model with *H*^1^ seminorm or convective regularisation, we are confident to state that these models are capable of estimating a meaningful source that can explain significantly more details of the observed signal.

In addition, we also evaluated (28) with 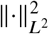 replaced by ║·║_*L*^∞^_, see Table 2 (right). Qualitatively, this leads to slightly different results for some datasets but still supports our main findings.

Let us illustrate the advantage of the convective regularisation on the basis of one particular dataset. In Fig. 10 we show the best result obtained for dataset number eight for three models. For this particular dataset, the model 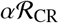 outperforms all other models according to Table 2.

**Fig. 10:**
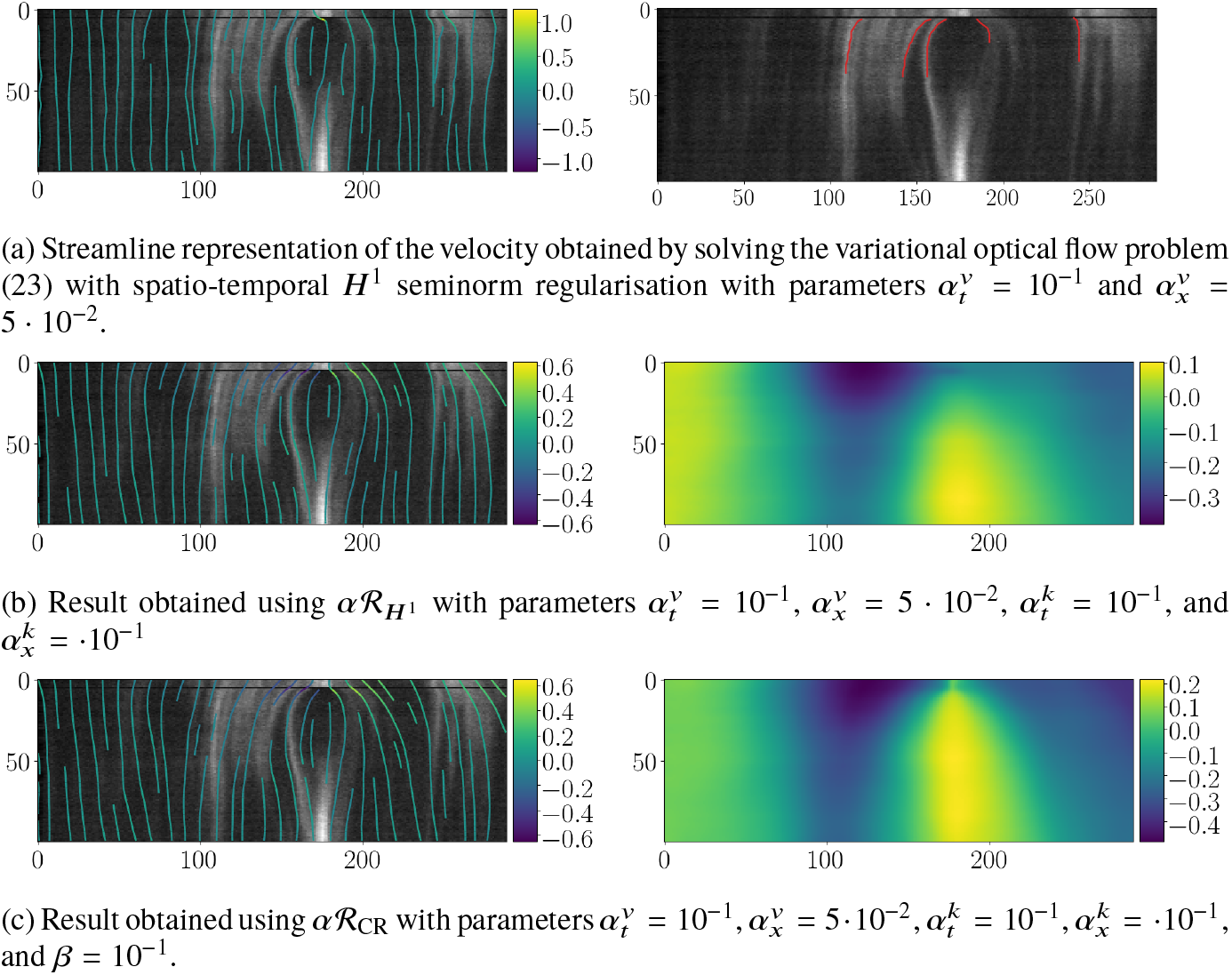
Best results obtained for dataset number eight in terms of the mean error (28) in the estimated velocity, see Table 2 (left).

Figure 10a (left) shows the best result for the optical flow model and Fig. 10a (right) the manually created tracks for this kymograph used to evaluate the computed velocities. Notice the inaccurate velocity between the cut ends shortly after the laser ablation. Moreover, towards the end of the sequence, where the both cut ends meet again, the characteristics seem inappropriate.

In Fig. 10b we display the result for the model 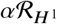. As can be seen in Fig. 10b (left), the estimated velocity is improved significantly in the cut region and in the problematic region towards the end of the sequence.

In Fig. 10c (left) we show the result for the model including the convective regularisation. i.e. for 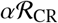. The estimated velocity appears visually as good as in the previous model with the additional advantage that it allows a larger magnitude shortly after the laser ablation. Observe that in both cases the estimated source is both spatially and temporally very regular, and apart from *beta* the same parameters 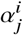 were selected. However, in Fig. 10c (right) the effect of the anisotropic regularisation is clearly visible in the cut region.

Moreover, in Fig. 11 we show the best obtained results for two other datasets. For comparison, the top row shows the same dataset as in Figs. 5 and 6.

**Fig. 11:**
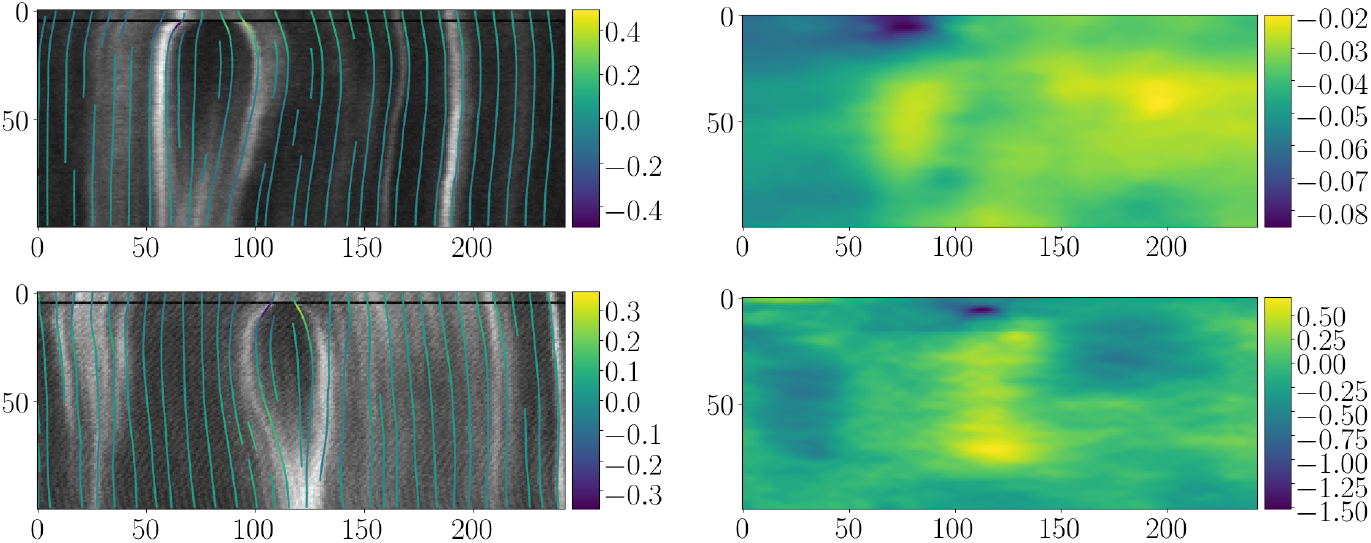
Best results obtained for dataset number three and five in terms of the mean error (28) in the estimated velocity, see Table 2 (left).

Finally, let us mention that, for the model 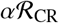, which we solve via Newton’s method, in total only 53 computations did not converge out of the 3125 parameter combinations tested on 15 different datasets.

### 4.5 Comparison based on a Mechanical Model of Tissue Formation

In Sec. 1 we have motivated the use of the non-homogenous continuity equation (1) mainly through mechanical models that are known to describe tissue formation, for example, in Drosophila. In order to see whether our variational formulation can reliably estimate velocity and source that both stem from such a process, we have implemented the partial differential equation-based model proposed in [26] to create synthetic data. In this model, the triplet (*m, v, σ*) solves the system

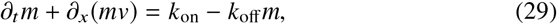

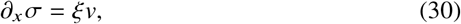

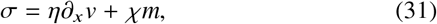

for (*t, x*) ∈ (0,*T*) × (0, 1), subject to the initial and boundary conditions

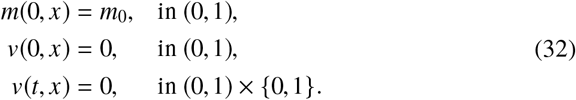

We refer to [45] for a derivation of this model.

Briefly, (29) is an advection–reaction equation modelling mass conservation of myosin molecules *m*, subject to rates of adsorption *k*_on_ > 0 and desorbtion *k*_off_ > 0, which we assume to be constants. The advection is determined by a mechanical problem, where mechanical balance (30) involves the stress *σ*(*t, x*) in the actin at the cell-cell junction and a friction force *ξv* exerted by the surrounding material, which we assume to be proportional to the velocity. Here, *ξ* > 0 is the coefficient of viscous drag. Finally, a constitutive model (31) of the junctional actin is proposed following e.g. [35, 23, 26] and involves viscous stresses and a pre-stress *χm* generated by myosin molecules. Again, *ηχ*, > 0 are constants. In addition, (32) enforces zero flux at the spatial boundaries, and *m*_0_ is an initial concentration. In our experiments we set it to

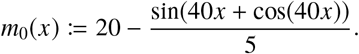

While such a model captures the essential features of actomyosin behaviour [35, 23, 47], its mean field approach means that away from the perturbation caused by the laser cut, the concentration *m* will equilibrate to the trivial solution (*m, v, σ*) = (*k*_on_ /*k*_off_, 0,*χk*_on_ /*k*_off_), which means that these parameters are uniform in space, and there will be no feature to track for an image analysis technique. This is also in contradiction with the experimental observations, where some material points along the cell-cell junctions exhibit accumulations of myosin that persist over time. One possible biophysical explanation for these accumulations is a locally larger density of actin binding sites.

This can be incorporated in the above model by introducing an additional variable *ρ*(*t, x*) and a constant 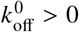 that modulates the off rate of myosin

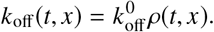

In addition, the density *ρ* obeys the conservation equation

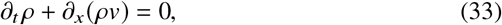

and satisfies an initial condition. In our experiments we set the initial *ρ* at *t* = 0 as

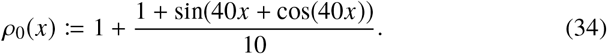

We solve the system (29)–(31) numerically, under additional consideration of (29), with a standard upwind finite volume discretisation paired with the forward Euler method. See, for instance, [46, Appx. B] for a brief description. For completeness, we briefly outline our implementation here.

We discretise the space-time domain [0,*T*]× [0, 1] using *N_t_* and *N_x_* equally spaced discretisation points in time and in space, respectively. For the discretisation of the unknowns we make use of a centred and a staggered grid, denoted by 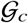 and 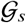, respectively. They are defined as

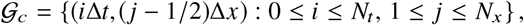

and as

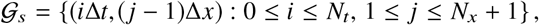

where Δ*t* = *T*/*N_t_* and Δ*x* = 1/*N_x_*. The concentrations and the stress are then discretised on the centred grid and the velocity on the staggered grid, leading to

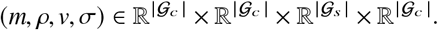

In further consequence, the finite volume discretisation, see e.g. [36, Chap. 4], of (29) reads

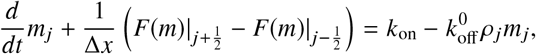

where 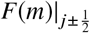 is the flux at the cell boundaries, that is, at the nodes of the staggered grid. Assuming that *m* is constant on each cell and using the upstream value of *m*, the flux 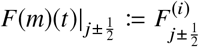 at time *t* = *i*Δ*t* at the boundaries can be written as

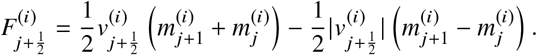

Similarly, we obtain 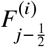 and, moreover, (17) results in zero flux at the boundaries. Then, approximating *d*/*dt* with forward finite differences yields

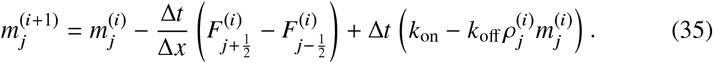

Analogously, an update equation for *ρ* is obtained.

The velocity at the current time step (*i*) is determined as follows. Using (30) to obtain *v* = *∂_x_σ*/*ξ* and substituting it into (31) yields the second-order elliptic partial differential equation

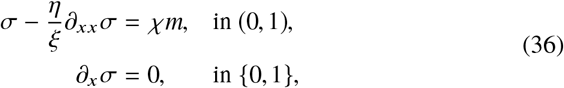

for the stress *σ*. Here, the zero Neumann boundary conditions follow from (32).

In each time step we solve (36), given the concentration *m*^(*i*)^ from the previous time instant, with a standard finite-difference scheme using centred differences on 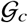. Then, with the help of (30) the velocity at nodes *j* + 1/2 can be approximated with

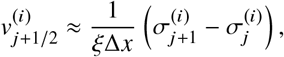

where we also use the fact that *v* is zero outside the spatial domain.

Finally, the concentration 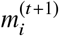 is updated according to (35) and *ρ* to its corresponding equation, and the procedure is repeated for the next time instant. The time interval is adjusted in each step so that the Courant–Friedrichs–Lewy condition is satisfied. This typically leads to intermediate results that are not recorded.

In our experiments we set *N_t_* = 300 and *N_x_* = 300, and the parameters controlling the time stepping were set to *T* = 0.1 and to Δ*t* = 2.5 · 10^−6^. The mechanical parameters in (30) and (31) were chosen as *η* = 1, *ξ* = 0.1, and as *χ* = 1. The parameters in (31) related to the source were set to *k*_on_ = 200 and to 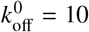.

In order to evaluate our approach described in Sec. 2.2, we conducted two experiments based on this mechanical model. In the first experiment, we used the solution method outlined above to generate a concentration *m*, which was then used as input to our variational formulation defined in (13). In the second experiment, we generated a concentration by solving (29) for a set velocity *v*, effectively removing the mechanical part of the model.

#### Unknown Velocities

In this experiment, we solved the mechanical model (29)–(31) together with (33) numerically as outlined above. In order to simulate a laser ablation, the concentration *m*_0_ is set to zero at nodes within the interval [0.495, 0.505]. In this way, a disruption (or loss) of concentration is simulated. Figure 12 (top) shows the solution (*m, v*) and the resulting source 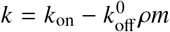.

**Fig. 12:**
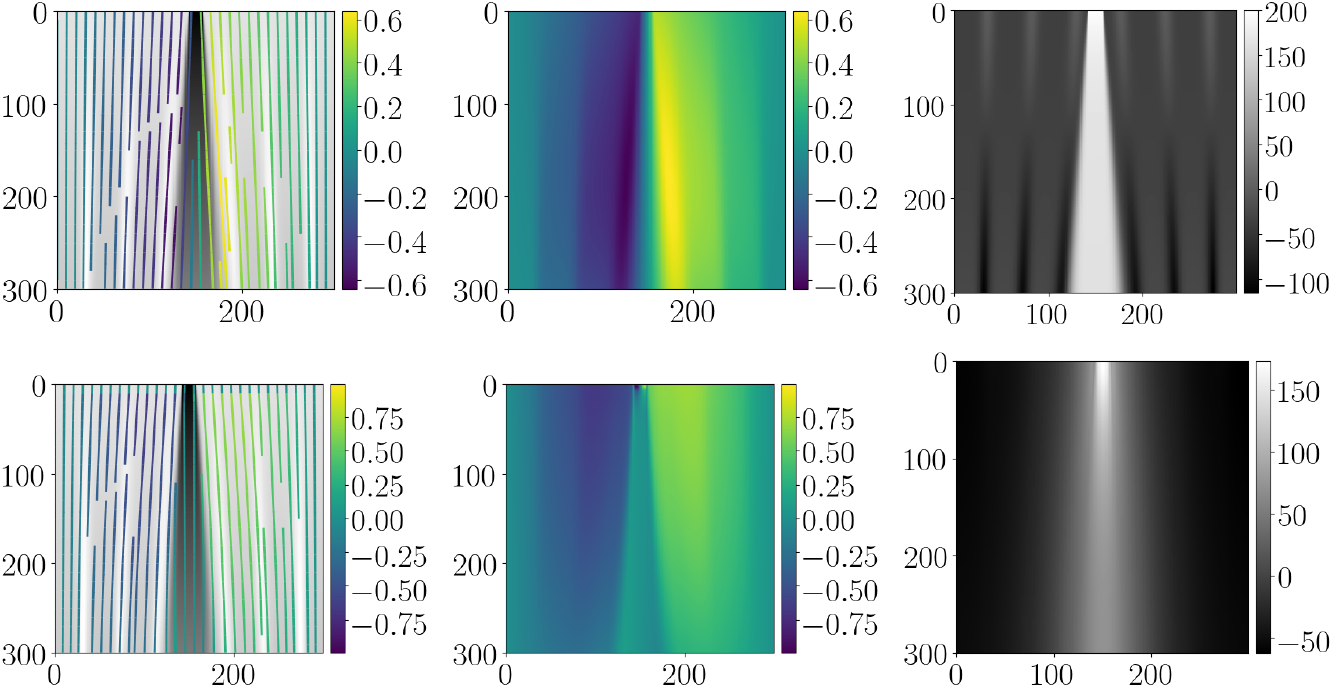
Shown is in the top row the solution (*m, v, k*) of the extended mechanical model (29)–(31). For simplicity, *ρ* and *σ* are omitted. The bottom images depict an approximate minimiser *w* = (*v, k*)^⊤^ obtained using (10) with 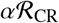. The left column shows the myosin concentration *m* together with streamlines obtained from the velocities, the middle column depicts the velocity *v*, and the right column illustrates the source *k*.

We then solved (13) with 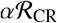 numerically based on the generated concentration. This is achieved by setting *f* := *m*. However, in order to match the boundary conditions in (32) we also used zero Dirichlet boundary conditions for *v* in (18) at *x* ∈ {0, 1} and at *t* = 0. In Fig. 12 (bottom), we depict an approximate minimiser. The parameters were set to 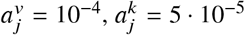, and to *β* = 10^−6^.

Observe that both the velocity and the source are estimated approximately and are within the correct order of magnitude. However, let us add that the estimated source appears quite regular in comparison to the simulation, even though the regularisation parameters 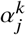 were set comparably small.

#### Predefined Velocities

In the next experiment, we removed the mechanical part and solved just (29) and (33) with a predefined velocity field *v* that could potentially resemble a laser nanoablation in cell membranes as pictured, for example, in Fig. 2.

We generated a velocity profile as follows. First, we define a characteristic *ϕ* : [0,*T*] → ℝ that is supposed to follow a cut end via the ordinary differential equation

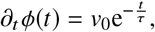

where the constant *v*_0_ > 0 is the initial velocity at time zero. Integrating with respect to time yields the integral curve

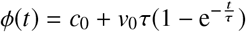

where *c*_0_ ≥ 0 is a constant defining the starting point of the curve. We then define a velocity field

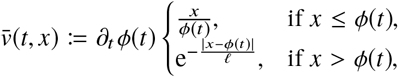

so that 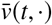 is linear in the interval [0, *ϕ*(*t*)] and decays exponentially in (*ϕ*(*t*), +∞). Here, *τ, ℓ* > 0 are constants and control the decay. Finally, we shift the origin to 1/2, reflect 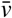, and obtain

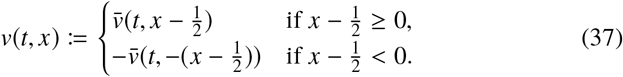

In Fig. 13 (top) we illustrate the solution (*m, v, k*) for this scenario, where *v* is set as in (37). The parameter *c*_0_ was set to *c*_0_ = 0.05 to match the width of the simulated laser ablation. The other parameters were set to *v*_0_ = 1, *τ* = 0.075, and to *ℓ* = 0.05. All other settings and parameters were kept as in the previous experiment.

**Fig. 13:**
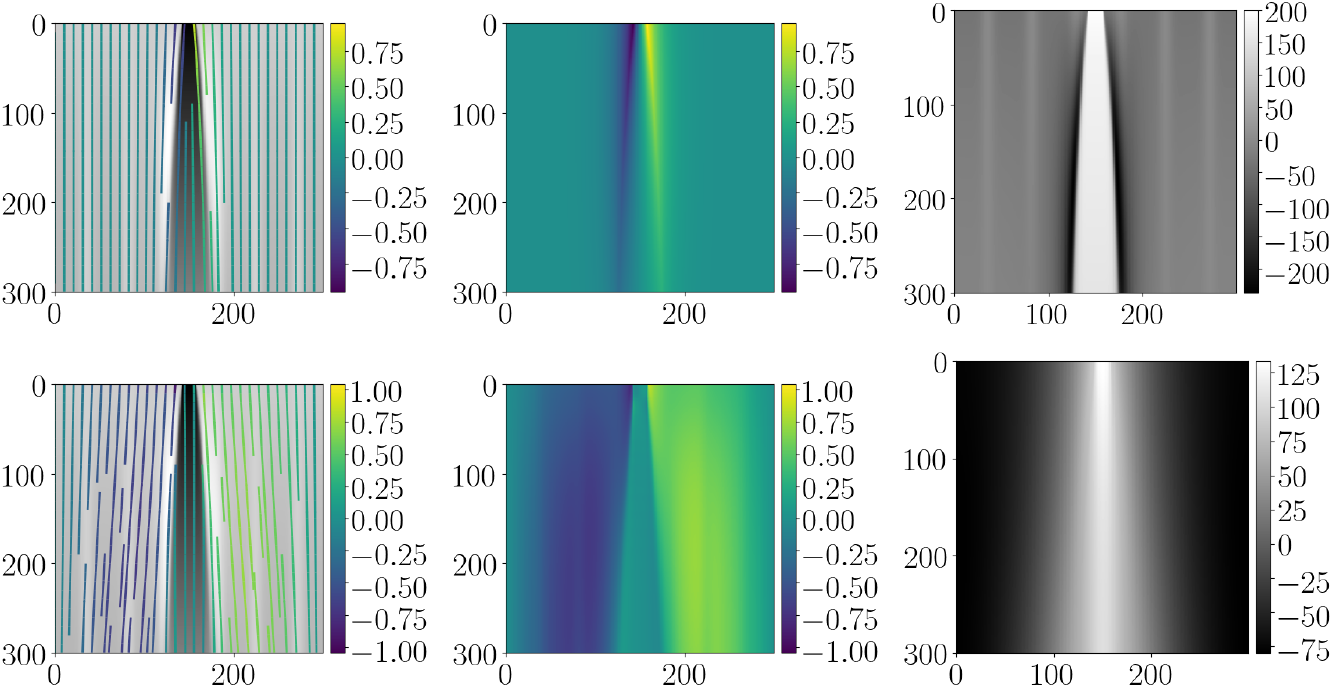
The top row shows functions (*m, k*) obtained by solving (29) and (33) given a set velocity *v*. The bottom images depict an approximate minimiser *w* = (*v, k*)^⊤^ obtained using (10) with 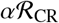. The left column shows the myosin concentration m together with streamlines obtained from the velocities, the middle column depicts the velocity *v*, and the right column illustrates the source *k*.

Then, we minimised (13) together with convective regularisation numerically based on the resulting concentration *m*. Figure 13 (bottom) shows the estimated velocity and source pair for the synthetic data. A similar behaviour as in the previous experiment can be observed.

## 5 Conclusions

In this article, we have investigated a variational model for joint velocity estimation and source identification in challenging fluorescence microscopy data of live Drosophila embryos that show the controlled destruction of tissue. We exploited the fact that a large proportion of tissue deformation occurs along one space dimension and allows to create kymographs. Our formulation is grounded on one-dimensional mechanical models of tissue formation and is based on the nonhomogenous continuity equation. We have discussed the ill-posedness of this problem and devised a well-posed variational formulation using convective regularisation of the source. Moreover, we have shown the connection of convective regularisation of the source to anisotropic diffusion. In a thorough experimental evaluation, we have demonstrated that motion estimation can benefit from simultaneously estimating a source and that convective regularisation may help to estimate velocities more accurately. Our numerical results show that this method could potentially help to quantify the reaction term in biological models of tissue formation. The extension of our models to more than one space dimension is left for future research.

## Acknowledgements

LFL and CBS acknowledge support from the Leverhulme Trust project “Breaking the non-convexity barrier”, the EPSRC grant EP/M00483X/1, the EPSRC Centre Nr. EP/N014588/1, the RISE projects ChiPS and NoMADS, the Cantab Capital Institute for the Mathematics of Information, and the Alan Turing Institute. ND and JE were supported by ANR-11-LABX-0030 “Tec21”, by a CNRS Momentum grant, and by IRS “AnisoTiss” of Idex Univ. Grenoble Alpes. ND and JE are members of GDR 3570 MecaBio and GDR 3070 CellTiss of CNRS. Some of the computations were performed using the Cactus cluster of the CIMENT infrastructure, supported by the Rhône-Alpes region (GRANT CPER07_13 CIRA) and the authors thank Philippe Beys, who manages the cluster. Overall laboratory work was supported by Wellcome Trust Investigator Awards to BS (099234/Z/12/Z and 207553/Z/17/Z). ES was also supported by a University of Cambridge Herchel Smith Fund Postdoctoral Fellowship. The authors also wish to thank Pierre Recho for fruitful discussions and the re-use of his numerical simulation code.

## Footnotes

1 https://doi.org/10.5281/zenodo.3263889

2 https://doi.org/10.5281/zenodo.3257654

## References

1. M. S. Alnæs, J. Blechta, J. Hake, A. Johansson, B. Kehlet, A. Logg, C. Richardson, J. Ring, M. E. Rognes, and G. N. Wells. The FEniCS project version 1.5. Archive of Numerical Software, 3(100):9–23, 2015.

2. F. Amat, E. W. Myers, and P. J. Keller. Fast and robust optical flow for time-lapse microscopy using super-voxels. Bioinformatics, 29(3):373–380, 2013.

3. A. A. Amini. A scalar function formulation for optical flow. In J.-O. Eklundh, editor, Proceedings of the 3rd European Conference on Computer Vision, volume 1, pages 123–131. Springer Berlin Heidelberg, 1994.

4. R. Andreev, O. Scherzer, and W. Zulehner. Simultaneous optical flow and source estimation: Space–time discretization and preconditioning. Appl. Numer. Math., 96:72–81, October 2015.

5. G. Aubert and P. Kornprobst. Mathematical problems in image processing, volume 147 of Applied Mathematical Sciences. Springer, New York, 2 edition, 2006. Partial differential equations and the calculus of variations, With a foreword by Olivier Faugeras.

6. D. Béréziat, I. Herlin, and L. Younes. A generalized optical flow constraint and its physical interpretation. In Proceedings of the IEEE Conference on Computer Vision and Pattern Recognition, volume 2, pages 487–492, 2000.

7. G. B. Blanchard, A. J. Kabla, N. L. Schultz, Lucy C Butler, B. Sanson, N. Gorfinkiel, L. Mahadevan, and R. J. Adams. Tissue tectonics: morphogenetic strain rates, cell shape change and intercalation. Nat. Meth., 6(6):458–464, may 2009.

8. J. T. Blankenship, S. T. Backovic, J. S. P. Sanny, O. Weitz, and J. A. Zallen. Multicellular rosette formation links planar cell polarity to tissue morphogenesis. Dev. Cell, 11(4):459–470, oct 2006.

9. A. Boquet-Pujadas, T. Lecomte, M. Manich, R. Thibeaux, E. Labruyère, N. Guillén, J.-C. Olivo-Marin, and A. C. Dufour. BioFlow: a non-invasive, image-based method to measure speed, pressure and forces inside living cells. Sci. Rep., 7(1), August 2017.

10. K. Boric, P. Orio, T. Viéville, and K. Whitlock. Quantitative analysis of cell migration using optical flow. PLOS ONE, 8:1–11, July 2013.

11. M. Burger, H. Dirks, and C.-B. Schönlieb. A variational model for joint motion estimation and image reconstruction. SIAM J. Imaging Sciences, 11(1):94–128, 2018.

12. A. R. Chaphalkar, K. Jain, M. S. Gangan, and C. A. Athale. Automated multi-peak tracking kymography (amtrak): A tool to quantify sub-cellular dynamics with sub-pixel accuracy. PLOS ONE, 11(12), 12 2016.

13. N. Chenouard, J. Buisson, I. Bloch, P. Bastin, and J. C. Olivo-Marin. Curvelet analysis of kymograph for tracking bi-directional particles in fluorescence microscopy images. In IEEE International Conference on Image Processing, pages 3657–3660, Sept 2010.

14. J.P. Cocquerez, L. Chanas, and J. Blanc-Talon. Simultaneous inpainting and motion estimation of highly degraded video-sequences. In J. Bigun and T. Gustavsson, editors, Image Analysis, volume 2749 of Lecture Notes in Computer Science, pages 685–692. Springer Berlin Heidelberg, 2003.

15. T. Corpetti, D. Heitz, G. Arroyo, É. Mémin, and A. Santa-Cruz. Fluid experimental flow estimation based on an optical-flow scheme. Expe. Fluids, 40(1):80–97, 2006.

16. T. Corpetti, É. Mémin, and P. Pérez. Dense estimation of fluid flows. IEEE Trans. Pattern Anal. Mach. Intell., 24(3):365–380, 2002.

17. R. Courant and D. Hilbert. Methods of mathematical physics. Vol. I. Interscience Publishers, Inc., New York, N.Y., 1953.

18. G. Crippa. The flow associated to weakly differentiable vector fields. PhD thesis, Classe di Scienze Matematiche, Fisiche e Naturali, Scuola Normale Superiore di Pisa / Institut für Mathematik, Universität Zürich, 2007.

19. B. Dacorogna. Direct methods in the calculus of variations, volume 78 of Applied Mathematical Sciences. Springer, New York, second edition, 2008.

20. M. Dawood, C. Brune, O. Schober, M. Schäfers, and K. P. Schäfers. A continuity equation based optical flow method for cardiac motion correction in 3D PET data. In H. Liao, P. J. Edwards, X. Pan, Y. Fan, and G.-Z. Yang, editors, Medical Imaging and Augmented Reality, volume 6326 of Lecture Notes in Computer Science, pages 88–97. Springer, Berlin, Heidelberg, 2010.

21. M. Drechsler, L. F. Lang, H. Dirks, M. Burger, C.-B. Schönlieb, and I. M. Palacios. Optical flow analysis reveals that kinesin-mediated advection impacts on the orientation of microtubules. bioRxiv, 2019.

22. S. J. England, G. B. Blanchard, L. Mahadevan, and R. J. Adams. A dynamic fate map of the forebrain shows how vertebrate eyes form and explains two causes of cyclopia. Development, 133(23):4613–4617, nov 2006.

23. J. Étienne, J. Fouchard, D. Mitrossilis, N. Bufi, P. Durand-Smet, and A. Asnacios. Cells as liquid motors: Mechanosensitivity emerges from collective dynamics of actomyosin cortex. 112(9):2740–2745, feb 2015.

24. L. C. Evans. Partial Differential Equations, volume 19 of Graduate Studies in Mathematics. American Mathematical Society, Providence, RI, second edition, 2010.

25. R. Fernandez-Gonzalez, S. de Matos Simoes, J.-C. Röper, S. Eaton, and J. A. Zallen. Myosin II dynamics are regulated by tension in intercalating cells. Dev. Cell, 17(5):736–743, nov 2009.

26. E. Hannezo, B. Dong, P. Recho, J.-F. Joanny, and S. Hayashi. Cortical instability drives periodic supracellular actin pattern formation in epithelial tubes. Proc. Nat. Acad. Sci. U.S.A., 112(28):8620–8625, June 2015.

27. H.W. Haussecker and D.J. Fleet. Computing optical flow with physical models of brightness variation. IEEE Trans. Pattern Anal. Mach. Intell., 23(6):661–673, June 2001.

28. C.-P. Heisenberg and Y. Bellaïche. Forces in tissue morphogenesis and patterning. Cell, 153(5):948–962, may 2013.

29. D. Heitz, E. Mémin, and Ch. Schnörr. Variational fluid flow measurements from image sequences: synopsis and perspectives. Expe. Fluids, 48(3):369–393, 2010.

30. B. K. P. Horn and B. G. Schunck. Determining optical flow. Artif. Intell., 17:185–203, 1981.

31. Y. Huang, L. Hao, H. Li, Z. Liu, and P. Wang. Quantitative analysis of intracellular motility based on optical flow model. J. Healthc. Eng., 2017:1–10, 2017.

32. J. Huisken and D. Y. R. Stainier. Selective plane illumination microscopy techniques in developmental biology. Development, 136(12):1963–1975, may 2009.

33. J. A. Iglesias and C. Kirisits. Convective regularization for optical flow. In M. Bergounioux, G. Peyre, C. Schnörr, J.B. Caillau, and T. Haberkorn, editors, Variational Methods in Imaging and Geometric Control, Radon Series on Computational and Applied Mathematics, pages 184–201. Walter de Gruyter GmbH & Co. KG, 2016.

34. C. Kirisits, L. F. Lang, and O. Scherzer. Optical flow on evolving surfaces with space and time regularisation. J. Math. Imaging Vision, 52(1):55–70, May 2015.

35. K. Kruse, J. F. Joanny, F. Jülicher, J. Prost, and K. Sekimoto. Generic theory of active polar gels: a paradigm for cytoskeletal dynamics. Eur. Phys. J. E, 16(1):5–16, jan 2005.

36. R. J. LeVeque. Finite Volume Methods for Hyperbolic Problems. Cambridge Texts in Applied Mathematics. Cambridge University Press, 2002.

37. C. M. Lye, G. B. Blanchard, H. W. Naylor, L. Muresan, J. Huisken, R. J. Adams, and B. Sanson. Mechanical coupling between endoderm invagination and axis extension in drosophila. PLOS Biol., 13(11), nov 2015.

38. P. Mangeol, B. Prevo, E. J. G. Peterman, and E. Holzbaur. KymographClear and KymographDirect: two tools for the automated quantitative analysis of molecular and cellular dynamics using kymographs. Mol. Biol. Cell, 27(12):1948–1957, 2016.

39. C. Melani, M. Campana, B. Lombardot, B. Rizzi, F. Veronesi, C. Zanella, P. Bourgine, K. Mikula, N. Peyriéras, and A. Sarti. Cells tracking in a live zebrafish embryo. In Proceedings of the 29th Annual International Conference of the IEEE Engineering in Medicine and Biology Society (EMBS 2007), pages 1631–1634, 2007.

40. B. Monier, A. Pélissier-Monier, A. H. Brand, and B. Sanson. An actomyosin-based barrier inhibits cell mixing at compartmental boundaries in drosophila embryos. Nat. Cell Biol., 12(1):60–65, dec 2010.

41. S. Neumann, R. Chassefeyre, G. E. Campbell, and S. E. Encalada. KymoAnalyzer: a software tool for the quantitative analysis of intracellular transport in neurons. Traffic, 18(1):71–88, 2016.

42. M. Nishikawa, S. R. Naganathan, F. Jülicher, and S. W. Grill. Controlling contractile instabilities in the actomyosin cortex. eLife, 6, jan 2017.

43. T. Preusser, M. Droske, C.S. Garbe, A. Telea, and M. Rumpf. A phase field method for joint denoising, edge detection, and motion estimation in image sequence processing. SIAM J. Appl. Math., 68(3):599–618, 2008.

44. P. Quelhas, A. M. Mendonça, and A. Campilho. Optical flow based arabidopsis thaliana root meristem cell division detection. In A. Campilho and M. Kamel, editors, Image Analysis and Recognition, volume 6112 of Lecture Notes in Computer Science, pages 217–226. Springer Berlin Heidelberg, 2010.

45. P. Recho, T. Putelat, and L. Truskinovsky. Mechanics of motility initiation and motility arrest in crawling cells. J. Mech. Phys. Solids, 84:469–505, nov 2015.

46. P. Recho, J. Ranft, and P. Marcq. One-dimensional collective migration of a proliferating cell monolayer. Soft Matter, 12(8):2381–2391, 2016.

47. A. Saha, M. Nishikawa, M. Behrndt, C.-P. Heisenberg, F. Jülicher, and S. W. Grill. Determining physical properties of the cell cortex. Biophys. J., 110(6):1421–1429, 2016.

48. E. Scarpa, C. Finet, G. B. Blanchard, and B. Sanson. Actomyosin-driven tension at compartmental boundaries orients cell division independently of cell geometry In Vivo. Dev. Cell, 47(6):727–740.e6, dec 2018.

49. J. Schindelin, I. Arganda-Carreras, E. Frise, V. Kaynig, M. Longair, T. Pietzsch, S. Preibisch, C. Rueden, S. Saalfeld, B. Schmid, J.-Y. Tinevez, D. J. White, V. Hartenstein, K. Eliceiri, P. Tomancak, and A. Cardona. Fiji: an open-source platform for biological-image analysis. Nature, 9(7):676–682, jul 2012.

50. B. Schmid, G. Shah, N. Scherf, M. Weber, K. Thierbach, C. Campos Pérez, I. Roeder, P. Aanstad, and J. Huisken. High-speed panoramic light-sheet microscopy reveals global endodermal cell dynamics. Nat. Commun., 4:2207, 2013.

51. Ch. Schnörr. Determining optical flow for irregular domains by minimizing quadratic functionals of a certain class. Int. J. Comput. Vision, 6:25–38, 1991.

52. B. G. Schunck. The motion constraint equation for optical flow. In Proceedings of the 7th International Conference on Pattern Recognition, pages 29–22, 1984.

53. S. M. Song and R. M. Leahy. Computation of 3-D velocity fields from 3-D cine CT images of a human heart. IEEE Trans. Med. Imag., 10(3):295–306, 1991.

54. J. Weickert. Anisotropic Diffusion in Image Processing. Teubner, Stuttgart, 1998. European Consortium for Mathematics in Industry.

55. J. Weickert, A. Bruhn, T. Brox, and N. Papenberg. A survey on variational optic flow methods for small displacements. In O. Scherzer, editor, Mathematical Models for Registration and Applications to Medical Imaging, volume 10 of Mathematics in Industry, pages 103–136. Springer, Berlin Heidelberg, 2006.

56. J. Weickert and Ch. Schnörr. A theoretical framework for convex regularizers in PDE-based computation of image motion. Int. J. Comput. Vision, 45(3):245–264, 2001.

57. J. Weickert and Ch. Schnörr. Variational optic flow computation with a spatio-temporal smoothness constraint. J. Math. Imaging Vision, 14:245–255, 2001.

58. D. Weiskopf and G. Erlebacher. Overview of flow visualization. In C. D. Hansen and C. R. Johnson, editors, The Visualization Handbook, pages 261–278. Elsevier, Amsterdam, 2005.

59. R. P. Wildes, A. M. Amabile, M. J. and Lanzillotto, and T.S. Leu. Recovering estimates of fluid flow from image sequence data. Comput. Vis. Image Underst., 80(2):246–266, 2000.

60. L. Younes. Shapes and Diffeomorphisms, volume 171 of Applied Mathematical Sciences. Springer-Verlag Berlin, 2010.

61. L. Zhou, C. Kambhamettu, and D. B. Goldgof. Fluid structure and motion analysis from multispectrum 2D cloud image sequences. In Proceedings of the IEEE Conference on Computer Vision and Pattern Recognition, volume 2, pages 744–751, 2000.

